# Functional disconnection of associative cortical areas predicts performance during BCI learning

**DOI:** 10.1101/487074

**Authors:** Marie-Constance Corsi, Mario Chavez, Denis Schwartz, Nathalie George, Laurent Hugueville, Ari E. Kahn, Sophie Dupont, Danielle S. Bassett, Fabrizio De Vico Fallani

**Affiliations:** Inria Paris, Aramis project-team, F-75013, Paris, France; Institut du Cerveau et de la Moelle Epinière, ICM, Inserm, U 1127, CNRS, UMR 7225, Sorbonne Université, F-75013, Paris, France; CNRS, UMR 7225, F-75013, Paris, France; Institut du Cerveau et de la Moelle Epinière, ICM, Inserm U 1127, CNRS UMR 7225, Sorbonne Université, Ecole Normale Supérieure, ENS, Centre MEG-EEG, F-75013, Paris, France; Department of Bioengineering, School of Engineering and Applied Science, University of Pennsylvania, Philadelphia, PA 19104, USA; Department of Neurology, Perelman School of Medicine, University of Pennsylvania, Philadelphia, PA 19104, USA; Department of Physics and Astronomy, College of Arts and Sciences, University of Pennsylvania, Philadelphia, PA 19104, USA; Department of Electrical and Systems Engineering, School of Engineering and Applied Science, University of Pennsylvania, Philadelphia, PA 19104, USA

## Abstract

Brain-computer interfaces (BCIs) have been largely developed to allow communication, control, and neuro-feedback in human beings. Despite their great potential, BCIs perform inconsistently across individuals and the neural processes that enable humans to achieve good control remain poorly understood. To address this question, we performed simultaneous high-density electroencephalographic (EEG) and magnetoencephalo-graphic (MEG) recordings in a motor imagery-based BCI training involving a group of healthy subjects. After reconstructing the signals at the cortical level, we showed that the reinforcement of motor-related activity during the BCI skill acquisition is paralleled by a progressive disconnection of associative areas which were not directly targeted during the experiments. Notably, these network connectivity changes reflected growing automaticity associated with BCI performance and predicted future learning rate. Altogether, our findings provide new insights into the large-scale cortical organizational mechanisms underlying BCI learning, which have implications for the improvement of this technology in a broad range of real-life applications.

## Introduction

Voluntarily modulating brain activity is a skill that can be learned by capitalizing on the feedback presented to the user. Such an ability is typically used in neurofeedback control to self-regulate putative neural substrates underlying a specific behavior, as well as in brain-machine interfaces, or brain-computer interfaces (BCIs) (1), to directly regulate external devices. Despite the potential impact, from elucidating brain-behavior relationships (2) to identifying new therapeutics for psychiatric (3) and neurological disorders (4; 5), both neurofeedback and BCIs face several challenges that affect their usability. This includes inter-subject variability, uncertain long-term effects, and the apparent failure of some individuals to achieve self-regulation (6). To tackle these issues, investigators have searched for better decoders of neural activity (7) as well as for psychological factors (8) and appropriate training regimens (9) that can influence the user’s performance. On the other hand, neuroplasticity is thought to be crucial for achieving effective control and this has motivated a deeper understanding of the neurophysiological mechanisms of neurofeedback and BCI learning (10). At small spatial scales, the role of cortico-striatal loops with the associated dopaminergic and glutamatergic synaptic organization has been demonstrated in human and animal studies suggesting the procedural nature of neurofeedback learning (11). At larger spatial scales, evidence supporting the involvement of distributed brain areas related to control, learning, and reward processing has been provided in fMRI-based neurofeedback experiments (12). Recently, a motor imagery (MI) BCI study based on ECoG recordings showed that successful learning was associated with a decreased activity in the dorsal premotor, prefrontal, and posterior parietal cortices (13). To date, however, the emergence of large-scale dynamic cortical network changes in relation with BCI learning has not been tested directly.

On the above-mentioned grounds, we hypothesized that BCI learning would result from a large-scale dynamic brain connectivity reorganization. More specifically, based on previous evidence documenting user’s transition from a deliberate mental strategy to nearly automatic execution (13; 10), we expected that the reinforcement of activity in the cortical areas targeted by the BCI would be accompanied by a progressive decrease of functional integration in regions associated with the cognitive processes of human learning (14). Furthermore, we hypothesized that the characteristics of such network changes would contain valuable information for the prediction of the BCI learning rate.

To test these predictions, we simultaneously recorded high-density EEG and MEG signals in a group of naive healthy subjects during a simple MI-based BCI training consisting of 4 sessions over 2 weeks. For both EEG and MEG, we derived cortical activity signals by performing source-reconstruction and we studied the longitudinal task-modulated changes in different frequency bands. Specifically, we evaluated the spatial extension of the activated cortical areas as well as the regional connectivity strength over time. Finally, we tested their relationships with learning as measured by the BCI performance (see *Materials and Methods* for details).

## Results

### Behavioral performance and BCI controlling features

The BCI task consisted of a 1D, two-target right-justified box task (15) in which subjects learned to control the vertical position of a cursor that moved from left to right on the screen. To hit the up-target, subjects performed a motor imagery-based hand grasping (MI condition) and to hit the down-target, they had to stay at rest (Rest condition). At the beginning of each experimental session, we identified the controlling EEG features among the electrodes over the contra-lateral motor area and within the standard *α* and *β* frequency ranges during a calibration phase (*SI Figure S2)*.

We found that the ability to control the BCI significantly increased across sessions (days) but not within sessions (hours) (*SI Figure S3)*. The session effect was also present when we averaged the BCI accuracy scores across the runs of each session (one-way ANOVA, *F*_3,57_ = 13.9, *p* = 6.56.10^−7^). Despite the expected high inter-subject variability (> 8.95%), 16 subjects out of 20 learned to control the BCI by the end of the training, with accuracy scores above the chance level of 57% (16) (Figure 1A, *SI Table S1)*.

We next investigated the characteristics of the EEG controlling features. From a spatial perspective, the electrodes above the primary motor area of the right hand (C3 and CP3) tended to better discriminate the MI and Rest mental states (Figure 1B). The most discriminant frequencies occurred between high-*α* and low-*β* ranges (Figure 1B). These results are in line with previous studies (17). Notably, we observed a progressive focus over CP3 and low-*β* ranges throughout the sessions.

Among the demographical and psychological items that we measured before the experiment (see *SI Materials and methods*), only the kinesthetic imagery score (18) moderately predicted the ability of subjects to control the BCI (Spearman, r = 0.45, p = 0.045). However, these items could not predict the evolution of BCI accuracy over time (*SI Table S2)*, suggesting that BCI skill acquisition is a complex process that could not be simply explained by a reduced number of scores taken from questionnaires.

**Figure 1:**
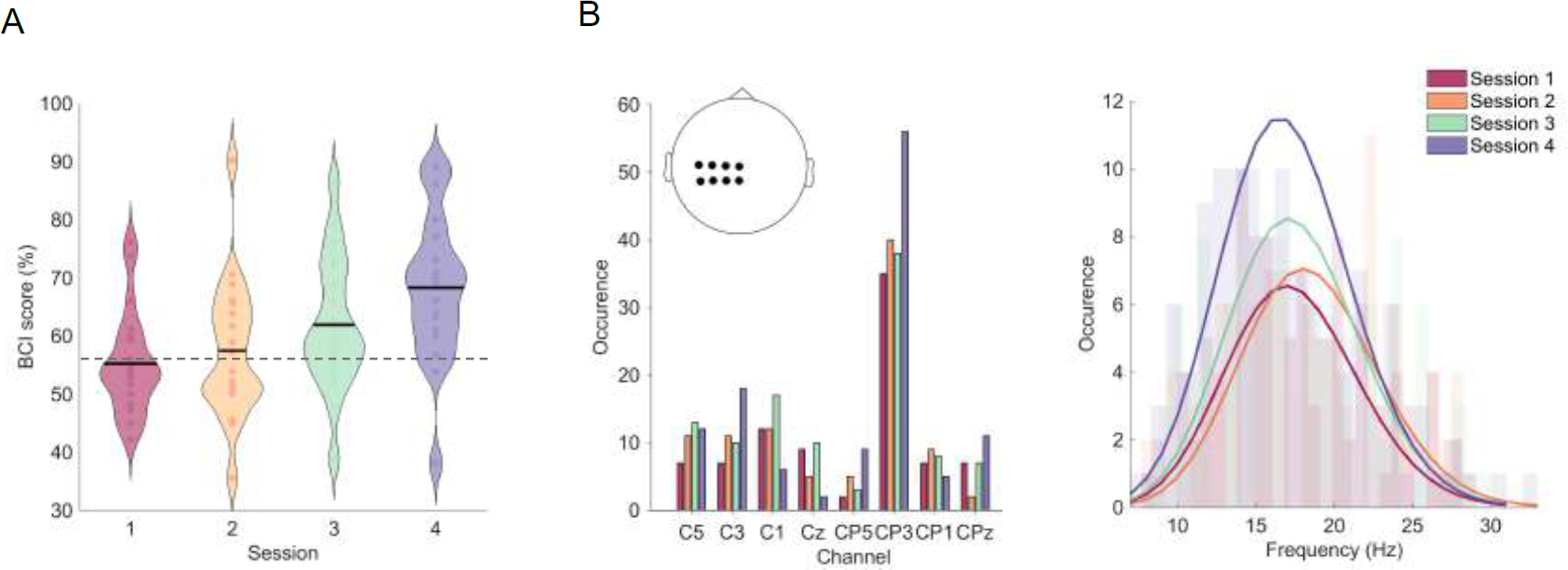
(A) Evolution of BCI performance over sessions. Individual performance is measured by considering the average BCI accuracy score (i.e. percentages of correctly hit targets) of the 96 trials in each session. In the violin plots, the black line corresponds to the group-averaged BCI score and the outer shape represents its distribution. The horizontal dashed grey line shows the chance level (57%), which is here considered as learning threshold. (B) Representation of the selected EEG controlling features across all subjects. On the left, we show occurrences obtained across subjects and sessions in terms of pre-selected channels; on the right, we show occurrences in terms of frequency bins selected over the sessions.

### Spatiotemporal cortical changes during training

We evaluated the spatiotemporal cortical changes associated with the BCI training by performing a statistical analysis of the MEG and EEG signals at the source-space level. Separately for each neuroimaging modality, we computed the associated task-related brain activity by statistically comparing the power spectra of the MI versus the Rest condition in each session, across subjects (*Materials and Methods*). In both *α* and *β* frequency ranges, we found a progressive involvement of distributed MEG sources in the cortical hemisphere contralateral to the movement (Figure 2). The involved regions exhibited a significant power decrease (p < 0.025), a phenomenon known as event-related desynchronization (ERD) which reflects sensorimotor brain activity (19).

**Figure 2:**
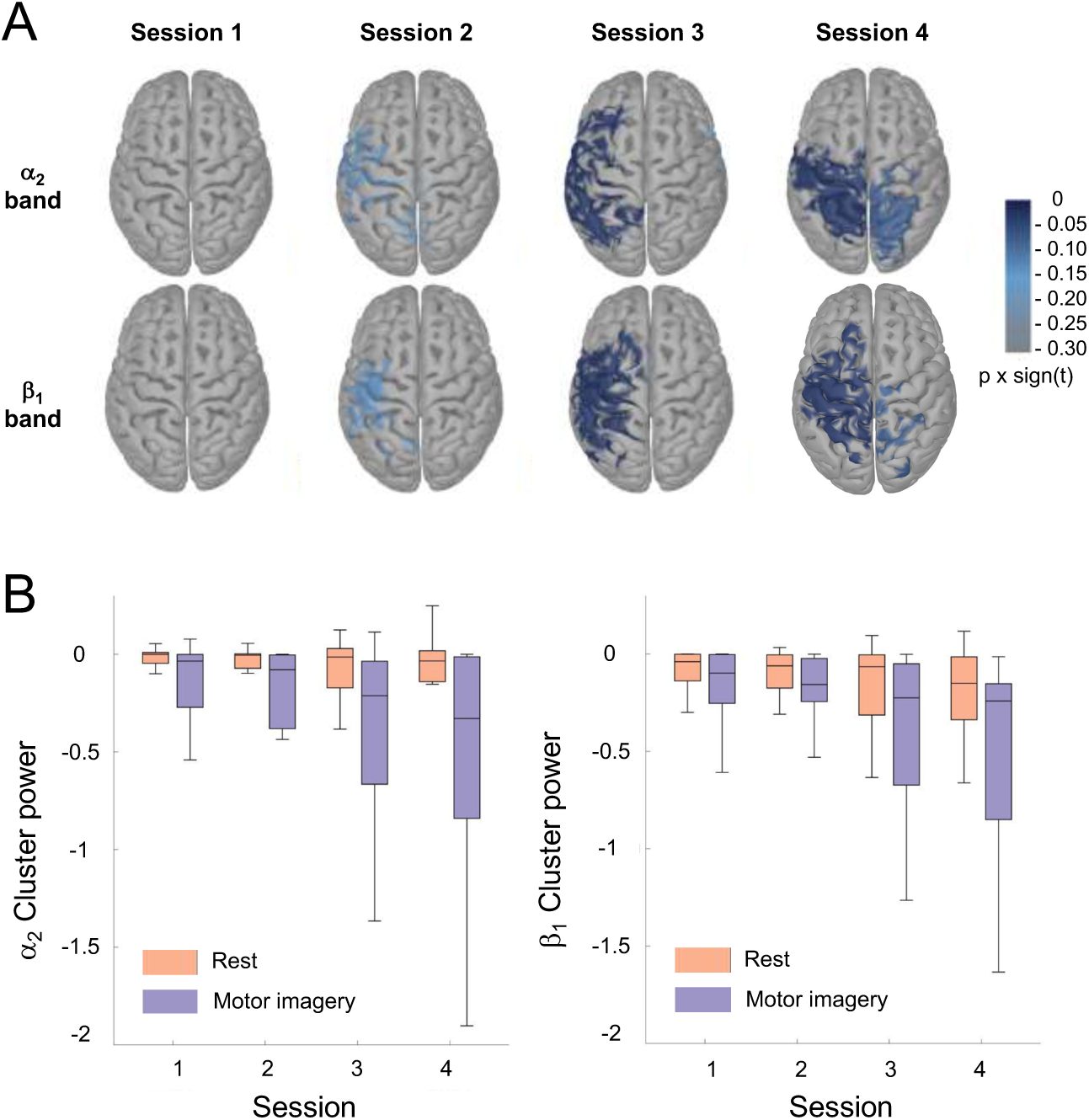
Cortical activity changes during BCI training. (A) Task-related activity maps obtained with MEG-source reconstructed power spectra in the *α*_2_ and *β*_1_ frequency band. The colors code the statistical difference obtained by contrasting motor-imagery and rest conditions through cluster-based permutation t-tests performed at the group level (*Material and methods*). For illustrative purposes, we show the obtained p-values multiplied by the sign of the t-values. (B) Normalized power spectra across the sessions for the motor-imagery (blue) and rest (red) condition. The significant clusters of activity in each individual were obtained with respect to the inter-stimulus intervals (ISI) (*Material and methods*). The group results for the *α*_2_ frequency band are illustrated by the boxplots on the left side of the panel, while results from the *β*_1_ frequency band are shown on the right. For illustrative purposes, we plotted the log transformed values. Similar results were obtained with EEG signals.

**Figure 3:**
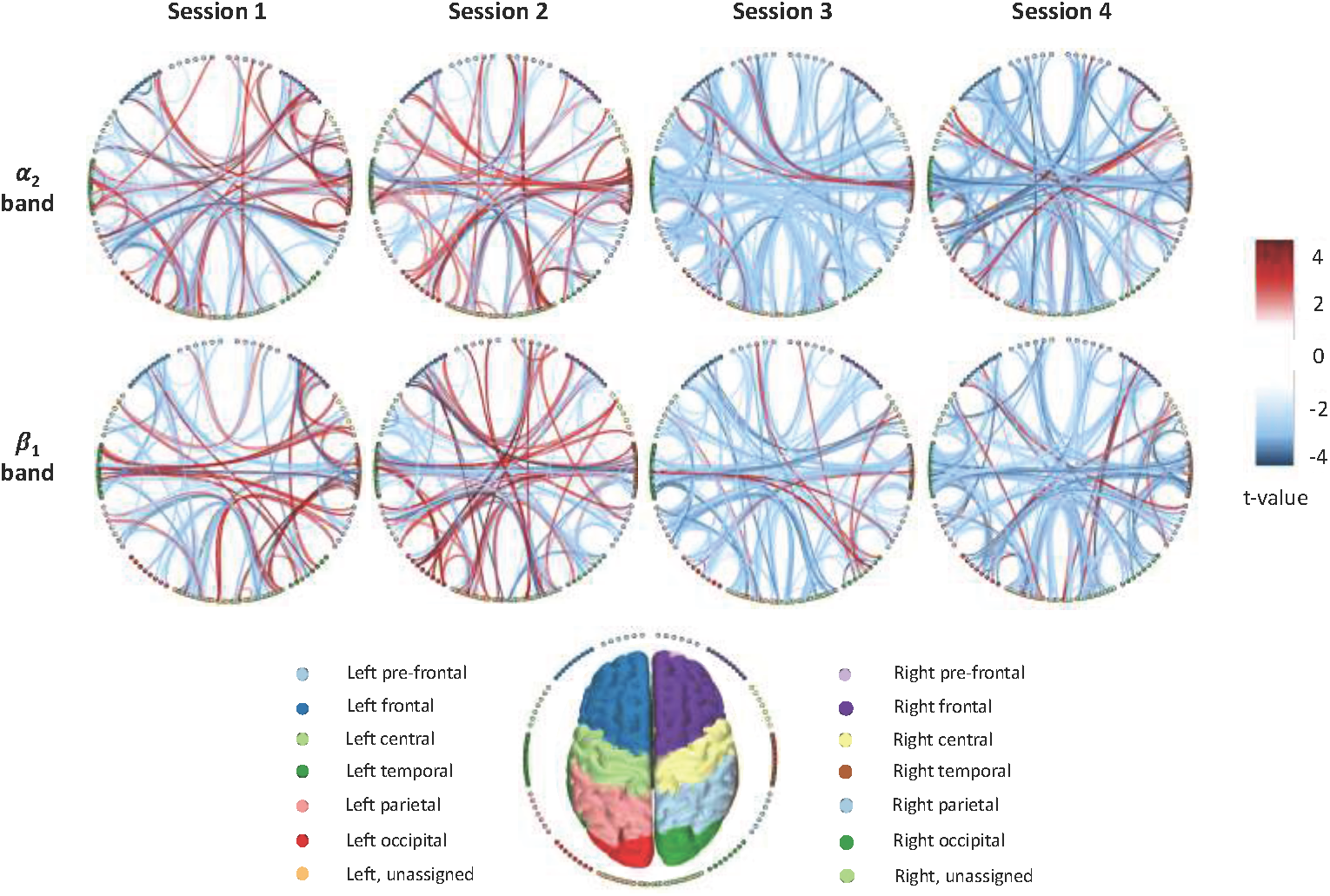
Cortical connectivity changes during BCI training. Task-related connectivity networks obtained with MEG-source reconstructed signals in the *α*_2_ and *β*_1_ band are represented on a circular graph. The nodes correspond to different regions of interest (ROIs) and the links code the statistical values resulting from a paired t-test performed between the motor-imagery and rest conditions performed at the group level (*Material and methods*). Only significant links (p < 0.005) are illustrated for the sake of simplicity. The color of each node, corresponds to a specific macro-area as provided by the Destrieux atlas. Similar results were obtained with EEG signals.

In session 3, ERDs were particularly significant in the *α*_2_ and *β*_1_ frequency bands, and mainly spanned the primary sensorimotor cortex (pre- and postcentral gyri, central sulcus, inferior and superior parts of the precentral sulcus) and secondary higher-order premotor and somatosensory areas (Figure 2, *SI Figures S5 and S8)*. At the end of training, ERDs were more localized in the contralateral paracentral lobule, precentral gyrus, and superior parietal lobule (Figure 2, *SI Figures S5 and S9)*, which are typically involved in hand motor tasks (20) as well as in motor imagery (21; 22) and motor learning (23). No other comparable significant differences were observed in the other frequency bands (*SI Figures S5)*.

To quantify these changes at the individual level, we calculated in each subject the size *C*_*S*_ and the relative power Δ_*P*_ of the most significant cluster of ERDs (*Materials and Methods*). These quantities exhibited a significant session effect only in the *α* and *β* frequency ranges (p < 0.03, *SI Table S3)*. Notably, BCI training was accompanied by larger (*C*_*S*_) and stronger (Δ_*P*_) clusters of task-related activity, including areas that are also outside the sensorimotor territory. These longitudinal changes were explained by the significant decrease of relative power in the MI condition (*F*_3,57_ = 4.82, *p* = 0.003; *F*_3,57_ = 3.09, *p* = 0.024 respectively in the *α*_2_ and *β*_1_ bands), while the Rest condition did not vary across sessions (*α*_2_) or varied with lower extent compared to MI (*β*_1_) (Figure 2). These findings confirm that during the training the subjects actually focused on how to perform MI of the hand.

Similar results, although with lower spatial resolution, were obtained when we considered EEG source-reconstructed signals (*SI Figures S4, S6 and S7)*. For the sake of simplicity, we will then focus on the results obtained with MEG while providing detailed EEG analysis in the supplementary materials.

### Functional connectivity and network analysis

To evaluate the cortical changes at the network level, we considered functional connectivity (FC) patterns that have been previously shown to be sensitive to BCI-related tasks (24) as well as to learning processes (25). For this purpose, we calculated the imaginary coherence between the source reconstructed signals of each pair of regions of interest (ROIs) corresponding to the Destrieux atlas (*Material and methods*). Imaginary coherence is a spectral measure of coherence weakly affected by volume conduction and spatial leakage (26; 27).

By statistically comparing the MEG-based FC values between MI and Rest conditions across subjects, we found a progressive decrease of task-related connectivity in both *α* and *β* frequency ranges along sessions (*SI Figures S11, S12)*. In *α* frequency ranges, the strongest decreases involved fronto-occipital (*α*_1_,*α*_2_) and parieto-occipital (*α*_2_) interactions. In *β* frequency ranges, significant decrements involved parieto-occipital (*β*_1_) but also fronto-central and bilateral temporal interactions (*β*_1_,*β*_2_).

For each subject, we quantified the regional connectivity changes by computing the relative node strength Δ_*N*_ in the *α* and *β* frequency ranges (*Materials and methods*). Significant across-session declines were spatially distributed involving bilaterally primary visual areas and associative regions (p < 0.025, *SI Figures S13)*. Specifically, in the *α*_2_ band, the Δ_*N*_ values of the ROIs typically associated with visuo-spatial attentional tasks (28) (middle occipital gyrus, cuneus) decreased significantly with the training (*SI Table S5)*. In the *β*_1_ band, we observed a significant reduction for the orbital part of the inferior frontal gyrus, which is involved in mental rotation (29) and working memory (30; 31) (*SI Table S5)*.

Collectively, the results indicate that BCI training is associated with a progressive reduction of integration among cortical systems that are specialized for different functions such as motor imagery and learning, visual attention, and working memory. Similar results from the analysis of EEG-based FC networks are reported in *SI Figures S10, S12, S13, SI Table S4*.

### Predictive neural markers of BCI performance

To better understand how the observed cortical changes in *α* and *β* frequency ranges were associated with performance, we next performed a repeated-measures correlation analysis which takes into account the longitudinal nature of our data (32). In general, we found strong correlations in the *α*_2_ and *β*_1_ frequency bands where both activity and connectivity metrics significantly correlated with the BCI performance within the same session (*SI Table S6, SI Figure S15)*. Higher BCI scores were associated with a higher number of task-related cortical sources *C*_*S*_ (Figure 4A) and with a stronger decrease of relative power Δ_*P*_ (Figure 4B). From a network perspective, better performance was associated with larger reduction of relative node strength Δ_*N*_ in associative areas and, to a minor extent, in primary visual regions (Figure 4C, *SI Figure S15)*.

**Figure 4:**
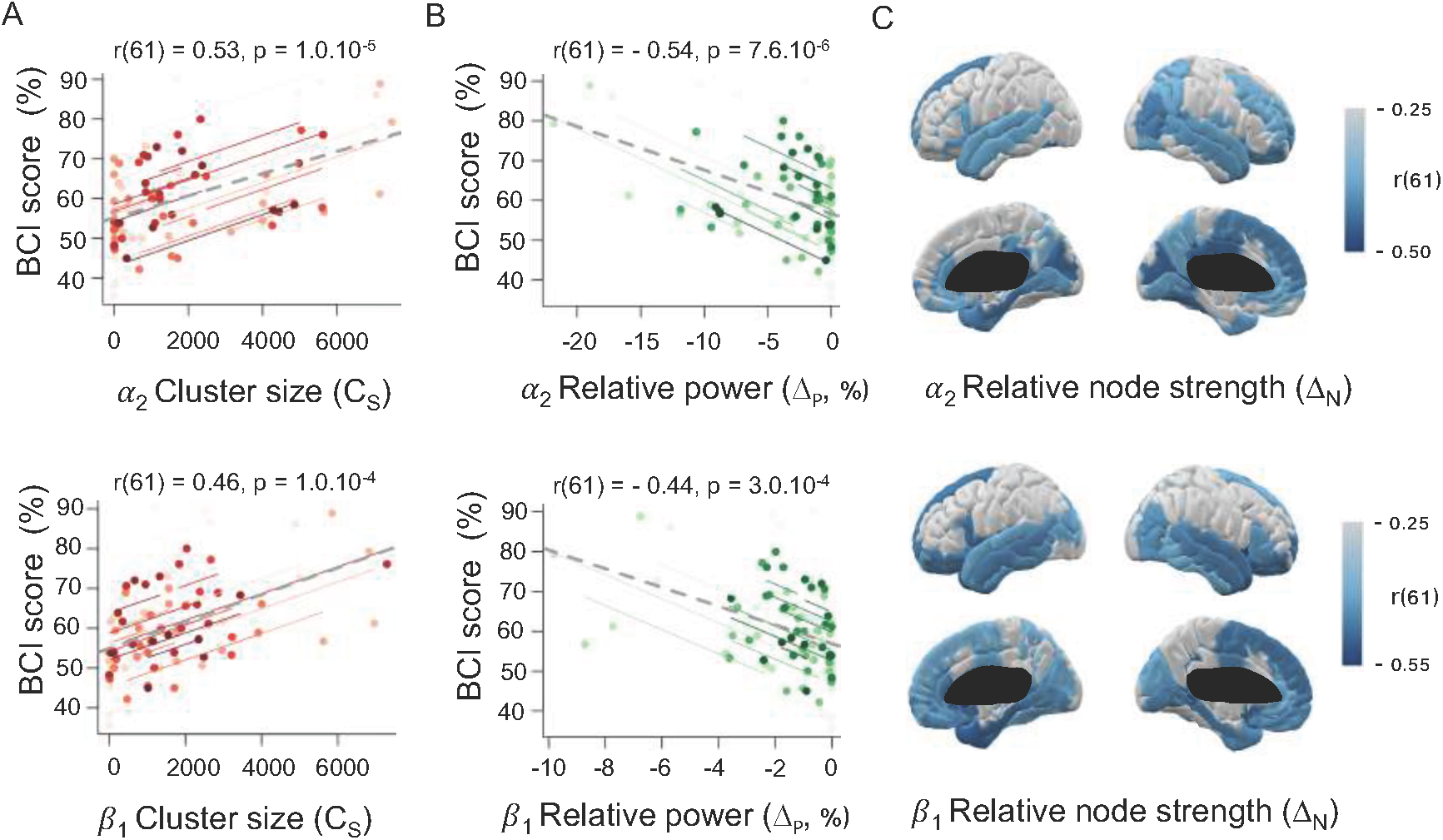
Correlation between activity/connectivity changes and BCI performance. The first row shows the results obtained in the *α*_2_ band. The second row shows the results obtained in the *β*_1_ band. (A) Scatter plots with cluster sizes *C*_*S*_ and the BCI scores of all the subjects. Colors identify the values obtained for the same individual across sessions. (B) Correlation values between the relative power Δ_*P*_ and the BCI scores of all the subjects. Same color conventions as in (A). (C) Correlation values between the relative node strengths Δ_*N*_ and the BCI scores in the same session. All correlations values (r) are calculated through a repeated-measures correlation coefficient, with a statistical threshold *α* = 0.05. For a detailed account of these results, see *SI Table S6* and *Figure S15*. Similar results were obtained with EEG signals.

Specifically, we observed significant correlations in regions known to be involved in cognitive aspects of human learning, such as decision making and memory consolidation (middle-anterior part of the cingulate gyrus) (33), change detection and shifts in behavior (posterior-ventral part of the cingulate gyrus) (34), as well as motion detection and tracking (lingual gyrus) (35; 36). Furthermore, we observed that areas known to be involved in both mental rotation and working memory (e.g. orbital part of the inferior frontal gyrus)(29; 30; 31), were also correlated with BCI scores. No other comparable significant differences were observed in the other frequency bands (*SI Table S6)*.

In terms of future prediction, we found that only the relative node strength (Δ_*N*_) was significantly correlated with the learning rate, defined as the relative difference of BCI accuracy between consecutive sessions (*p <* 0.05, *SI Figure S15, SI Table S6)*. Strongest correlations were found in *α*_2_ and *β*_1_, where higher values of Δ_*N*_ were associated with the a larger learning amount in the following session.

In these bands, the most predictive cortical areas were the anterior part of the cingulate gyrus and the orbital part of the inferior frontal gyrus (Figure 5), both known to be involved in human learning (37). Significant predictions were also reported for the fronto-marginal gyrus in the *β*_1_ band and for the the superior parietal lobule in the *α*_2_ band (*SI Figure S15)*, which are typically associated with learning and motor imagery tasks (38; 39; 21).

We obtained similar results from the analysis of EEG source-reconstructed activity and connectivity *SI Table S6, SI Figure S14*. Altogether, these findings demonstrate that the observed dynamic cortical changes at the network level were intrinsically associated with successful BCI learning.

**Figure 5:**
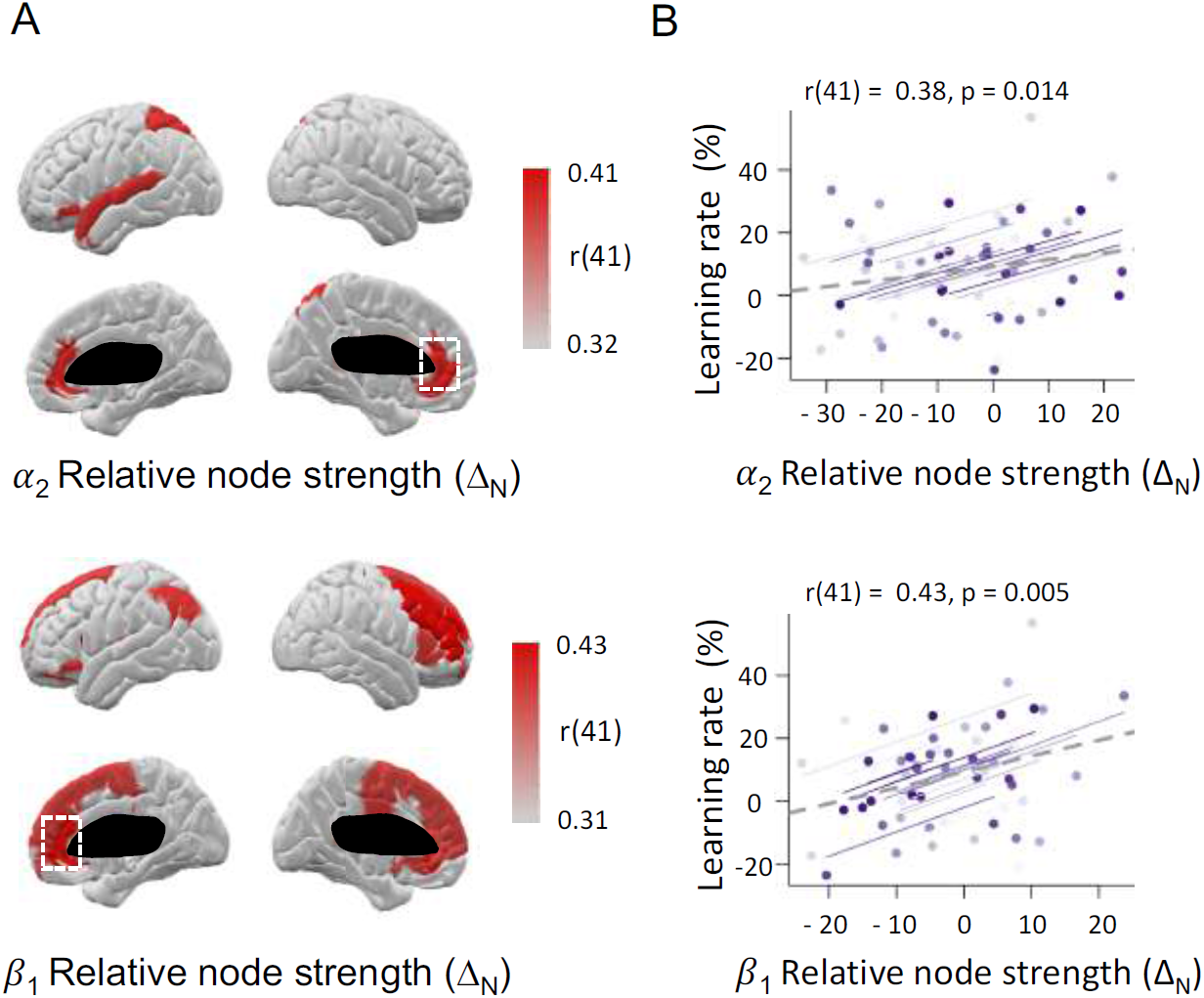
Prediction of BCI learning rate from regional connectivity strengths. The first row shows the results obtained in the *α*_2_ band. The second row shows the results obtained in the *β*_1_ band. (A) Colors show the correlation values for the ROIs with a significant effect p < 0.05. (B) Scatter plots show the values of relative node strengths Δ_*N*_ and the learning rates of all the subjects for the most significant ROIs (p < 0.002). Colors identify the values obtained for the same individual across sessions. The r values correspond to the repeated measures correlation coefficients. For a detailed account of these results, see *SI Figure S15*. Similar results were obtained with EEG signals.

## Discussion

### Neuroplasticity and motor learning

Identifying the large-scale neural mechanisms underlying plasticity is fundamental to understand human learning (23; 40; 41; 42). The ability to voluntarily modulate neural activity to control a BCI appears to be a learned skill. Investigators have repeatedly documented that task performance typically increases over the course of practice (43; 44), while BCI users often report transitioning from a deliberate cognitive strategy (e.g. MI) to a nearly automatic goal-directed approach focused directly on effector control. This evidence is indicative of a learning process taking place in the brain that is consistent with procedural motor learning. Efforts in understanding the neural dynamics underlying BCI skill acquisition have been made by using neuroimaging techniques in primates (43; 45) as well as in humans (46; 13). These works suggest that even if the use of a BCI only requires the modulation of activity in a motor-related brain area, a dynamic and distributed network of remote cortical areas is involved in the early acquisition of BCI proficiency. However, how such network reorganizes over time and which are the connectivity mechanisms subserving BCI learning is still largely unknown.

Here, we showed that a MI-based BCI learning is associated with a progressive decrease of functional integration of associative cortical regions and with the reinforcement of sensorimotor activity targeted by the experiment. Together with sensorimotor areas, associative areas play a crucial role in motor sequence learning as well as in abstract task learning (23; 40; 41; 42). In particular, we showed that MI-based BCI learning is accompanied by the disconnection of specific areas related to working memory and decision-making (Figure 3, *SI Figures S12-S13)*. Both these cognitive processes are known to be involved in the supervisory attentional system (47; 48), which is an important prerequisite for successful motor learning (49; 50; 51; 52) and also present during MI tasks (53). Altogether, we speculate that the observed progressive disconnection of associative areas would mainly reflect the attentional effort fading related to the optimization of the MI strategy to control the BCI and to the gradual skill acquisition.

### Network predictors of learning rate

Forecasting behavior from brain functioning is one of the main challenges in human neuroscience. In BCI contexts, the identification of neural features predicting BCI performance will allow to better design adaptive BCIs. Investigators have recognized the need for adaptive BCI architectures that accommodate the dynamic nature of the neural features used as inputs (54). Initial efforts have taken into account psychological (e.g. anxiety) or demographical items to predict BCI performance, but the associated results seem to be contradictory (55; 56; 57). Recently, FC-based metrics have been shown to correlate with the user’s performance suggesting potential strategies for improving MI-based BCI accuracy (58).

However, these findings only referred to the same experimental session and did not inform on the prediction in the future sessions. In this field, there is a critical need for biologically informed computational approaches to identify the neural mechanisms of BCI learning that predict future performance, thereby enabling the generalization of these results across subject cohorts, and the optimization of BCI architectures for individual users (59). Here, we showed that the regional connectivity strength of specific associative cortical areas is not only able to predict the BCI performance in the same session (Figure 4C) but can also predict the learning rate in the subsequent session (Figure 5). Notably, higher values of relative node strength Δ_*N*_ were associated with larger learning rates, indicating that the potential to improve performance is higher when the functional disconnection of the associative areas has not yet started. These findings could be used to inform future decisions on how to train individuals depending on the current properties of the functional brain network organization.

### BCI “illiteracy” and performance assessment

BCIs are increasingly used for control and communication as well as for the treatment of neurological diseases (60). While performance usually reaches high levels of accuracy (around 90 %), a non-negligible portion of users (between 15 % and 30 %) exhibit an inability to communicate with a BCI (61). This is a well-known phenomenon that is informally referred as to “BCI illiteracy”. Critically, it affects the usability of BCIs in the user’s daily life but the reasons (62; 63) and even the definition for such inability (64) is still under debate. On the one hand, different approaches have been proposed to solve the problem by improving feature extraction and decoding algorithms (54), combining different modalities (65), or taking into account the user’s profile (63). On the other hand, alternative accuracy metrics based on the separability of brain features, rather than on the simple count of successful control, have been shown to be more relevant for the proof-of-existence of subject’s learning (66).

Our results show that a functional reorganization of the cortical activity is taking place during BCI training. These findings suggest that BCI illiteracy could be an epiphenomenon biased by the nature of standard performance metrics which are affected by decoder recalibration (67), re-parameterizations of the BCI, and the application and adoption of better mental strategies (68; 59), among other factors. It is possible that other metrics, integrating both real performance and functional brain changes should be taken into account to better assess individual learning (66).

### Limitations

The temporal window of two weeks considered in our experiment prevents us from observing behavioral and neural changes over longer timescales and therefore, might not be sufficient to observe the full learning process (69). Here, BCI skill acquisition was paralleled by a progressive focused activity over the sensorimotor areas together with a loss of large-scale connectivity, which altogether indicate the initiation of an automaticity process typical of procedural motor learning (14). Future studies are necessary to assess whether and how the observed cortical patterns will evolve with longer BCI training.

While the BCI accuracy was highly variable across individuals, the group-averaged performance was relatively low as compared to the state-of-the-art (6). It is important to mention that the main goal of the present work was not to maximize the performance but to study the neural mechanisms underlying BCI learning. In this respect, all our experimental subjects were BCI-naive and exhibited on average an increase of performance reflecting a successful BCI skill acquisition.

From this work alone, we are unable to determine whether or not learning is the only possible modulator of the observed cortical changes. While no correlation has been found with behavioral factors (i.e. anxiety), complementary experiments could be designed to test whether the observed cortical changes are also modulated by fatigue or exogenous stimulants to increase motor excitability (i.e., transcranial direct current stimulation, tDCS (70)).

## Conclusion

Consistent with our hypothesis, we have identified specific cortical network changes that characterize dynamic brain reorganization during BCI training. We found that the progressive functional disconnection of associative areas is crucial for the BCI skill acquisition process. These network signatures varied over individuals and, more importantly, were significant predictors of the BCI learning rate. Taken together, our results offer new insights into the crucial role of brain network reconfiguration in the prediction of human learning.

## Materials and Methods

### Participants and experiment

Twenty healthy subjects (aged 27.5 ± 4.0 years, 12 men), all right-handed, participated in the study. Subjects were enrolled in a longitudinal EEG-based BCI training (twice a week for two weeks, *SI text)*. All subjects were BCI-naive and none presented with medical or psychological disorders. According to the declaration of Helsinki, written informed consent was obtained from subjects after explanation of the study, which was approved by the ethical committee CPP-IDF-VI of Paris. All participants received financial compensation at the end of their participation. The BCI task consisted of a standard 1D, two-target box task (15) in which the subjects modulated their *α* [8-12 Hz] and/or *β* [14-29 Hz] activity to control the vertical position of a cursor moving with constant velocity from the left to the right side of the screen. To hit the up-target, the subjects performed a sustained motor imagery of right-hand grasping (MI condition) and to hit the down-target they remained at rest (Rest condition). Each trial lasted 7 s and consisted of a 1 s of inter-stimulus, followed by 2 s of target presentation, 3 s of feedback and 1 s of result presentation. BCI control features (EEG electrode and frequency) were selected in a calibration phase at the beginning of each session, by instructing the subjects to perform the BCI tasks without any visual feedback (*SI Text, SI Figure 2)*. Experiments were conducted with a 74 EEG-channel system, with Ag/AgCl sensors (Easycap, Germany) placed according to the standard 10-10 montage. EEG signals were referenced to mastoid signals, with the ground electrode located at the left scapula, and impedances were kept lower than 20 kOhms. A system composed by 102 magnetometers and 204 gradiometers collected MEG data (Elekta Neuromag TRIUX MEG system). EEG and MEG signals were simultaneously recorded in a magnetic shielded room with a sampling frequency of 1 kHz and a bandwidth of 0.01-300 Hz. To digitize the head positions, we used the Polhemus Fastrak digitizer (Polhemus, Colchester, VT). Nasion, left and right pre-auricular points were used as landmark points to provide co-registration with the anatomical MRI. Four Head Position Indicator (HPI) coils were attached to the EEG cap. The subjects were seated in front of a screen at a distance of 90 cm. To ensure the stability of the position of the hands, the subjects rested their arms on a comfortable support, with palms facing upward. We also recorded electromyogram (EMG) signals from the left and right arm of the subjects. Expert bioengineers visually inspected EMG activity to ensure that subjects were not moving their forearms during the recording sessions. We carried out BCI sessions with EEG signals transmitted to the BCI2000 toolbox (71) via the Fieldtrip buffer (72).

### M/EEG preprocessing and source reconstruction

MEG signals were first preprocessed by applying the temporal extension of the Signal Space Separation (tSSS) to remove environmental noise with MaxFilter (73). MEG and EEG signals were downsampled to 250 Hz before performing an ICA (Independent Components Analysis) with the Infomax approach using the Fieldtrip toolbox (74; 72). The number of computed components corresponds to the number of channels, i.e. 72 for EEG (T9 and T10 were removed). Only the independent components (ICs) that contain ocular or cardiac artifacts were removed. The selection of the components was performed via a visual inspection of the signals (from both time series and topographies). On average, 2 ICs were removed. Data were then segmented into epochs of seven seconds corresponding to the trial period. Our quality check was based on the variance and the visual inspection of the signals. For each channel and each trial, we plotted the associated variance values. We kept a ratio below 3 between the noisiest and the cleanest trials. The percentage of removed trials was kept below 10 % of the total number of trials (75).

After having average referenced the signals, we performed source reconstruction by computing the individual head model with the Boundary Element Method (BEM) (76; 77). BEM surfaces were obtained from three layers associated with the subject’s MRI (scalp, inner skull, outer skull) that contain 1922 vertices each. Then, we estimated the sources with the weighted Minimum Norm Estimate (wMNE) (78; 79; 80) using the Brainstorm toolbox (81). Here, we used the identity matrix as the noise covariance matrix. The minimum norm estimate corresponds in our case to the current density map. We constrained the dipole orientations normal to the cortex. To perform the group analysis, we projected the sources estimated on each subject, and each session, onto the common template anatomy MNI-ICBM152 (82) via Shepard’s interpolation. From these estimated signals, we computed the associated power spectra. To identify the anatomical structures associated with the obtained clusters without restricting our work on motor or sensorimotor areas, we used the Destrieux atlas (83).

### Metrics and statistics

To take into account the subjects’ specificity (84), we used a definition of the *α* and *β* bands that rely on the Individual Alpha Frequency (IAF) obtained from a resting-state recording that lasted 3 minutes (with the subjects’ eyes open). Similarly to (4), the IAF corresponds to the first peak obtained between 6 and 12 Hz. The *α*_1_ ranges from IAF – 2 Hz to IAF, *α*_2_ from IAF to IAF + 2 Hz, *β*_1_ from IAF + 2 Hz to IAF + 11 Hz and *β*_2_ from IAF + 11 Hz to IAF + 20 Hz. For each subject, session, and trial in the source space, we computed the power spectra. We used the Welch method with a window length of 1 s and a window overlap ratio of 50 % applied during the feedback period that ranged from t = 3 s to t = 6 s. In the case of the group analysis presented in Figure 2A, we worked within the ICBM152-MNI template. Elsewhere, we used the individual anatomical space. To perform the analysis presented in Figure 2B, we computed statistical differences among activations recorded in the MI and the rest conditions at the group level or at the subject level via a paired t-test. Since we expected a desynchronization between the two conditions, we applied a one-tailed t-test. Statistics were corrected for multiple comparisons using the cluster approach (72; 81). We fixed the statistical threshold to 0.05, a minimum number of neighbors of 2 and a number of randomization of 500. Clustering was performed on the basis of spatial adjacency. Cluster-level statistics were obtained by using the sum of the t-values within every cluster.

To obtain the relative power Δ_*P*_, we computed the relative difference, in terms of power spectra, between the two conditions, as follows: 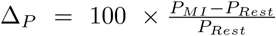, where *P*_*MI*_ and *P*_*Rest*_ correspond, respectively to the averaged power calculated across the cluster from MI and Rest trials. The cluster size *C*_*S*_ was obtained by estimating the number of elements that belong to the cluster that presented the best discrimination between the conditions. To perform the study for each condition separately (Figure 2), we normalized the power spectra with respect to the inter-stimulus interval (ISI) with the Hilbert transform, similar to the approach reported in (13). The connectivity analysis (Figure 3) was based on the cross-spectral estimation computed with the Welch method. To reduce the dimensionality, we extracted the first principal component obtained from the power spectra calculated across the dipoles within each ROI. Then, we computed the imaginary coherence between each pair of ROIs based on the definition proposed in (27). From the resulting connectivity matrix, we next computed the relative node strength Δ_*N*_ similarly to what we did for the relative power. The strength of the i-th node was here calculated by summing the values of the i-th row of the connectivity matrix. To evaluate the session effect on Δ_*P*_, *C*_*S*_, and Δ_*N*_ one-way repeated ANOVAs were applied with the session number as the intra-subject factor. In the specific case of Δ_*N*_, the ANOVA was performed separately for each ROI.

To estimate the correlations between BCI scores and, respectively, Δ_*P*_, *C*_*S*_, and Δ_*N*_,, we performed repeated-measures correlations (32) which control for non-independence of observations obtained within each subject without averaging or aggregating data. In the specific case of Δ_*N*_, the correlation analysis was performed separately for each ROI.

## Data availability

The data that support the findings of this study are available from the corresponding authors upon reasonable request.

## Acknowledgments

This work was partially supported by French program “Investissements d’avenir” ANR-10-IAIHU-06; “ANR-NIH CRCNS” ANR-15-NEUC-0006-02 and by NICHD 1R01HD086888-01. The funders had no role in study design, data collection and analysis, decision to publish, or preparation of the manuscript. This work was performed on a platform of France Life Imaging network partly funded by the grant “ANR-11-INBS-0006”.

## Authors contributions

MC, DS, NG, LH, SD, DSB and FDVF initiated research; MCC, MC, DS, LH, DSB and FDVF designed research; MCC, DS and LH performed research; MCC, DS, LH and AEK contributed analytic tools; MCC and AEK analyzed data; and MCC, DSB, and FDVF wrote the paper. All authors revised and approved the manuscript.

## Additional Information

Supplementary Information accompanies this paper.

