## Supplementary materials for "Functional disconnection of associative cortical areas predicts performance during BCI learning"

April 26, 2019

### Supplementary Materials

#### Supplementary text

##### Materials and Methods: BCI protocol

In order to study the neural mechanisms associated with the MI-based BCI learning, the subjects performed four sessions, twice a week, for two weeks. We used the one-dimensional, two-target, right-justified box task (1), where subjects performed a sustained MI (grasping) of the right hand to hit up-targets while remaining at rest to hit down-targets. Each run consisted of 32 trials with up and down targets, consisting in a grey vertical bar displayed on the right part of the screen, equally and randomly distributed across trials (see Figure S1). The experiment was divided into two phases:

1. The training phase consisted of five consecutive runs without any feedback. For a given trial, the first second corresponded to the inter-stimulus interval (ISI), while the target was presented during the subsequent five seconds. From the data obtained during this phase, contrast maps were computed to elicit the features, i.e. the (channel; frequency) couples of interest that best discriminate the subjects' mental state over the left motor area (see Figure S2). For that purpose, we used the R-square as a metric of such a discrimination between the conditions (2)).

2. The testing phase consisted of six runs with a cursor feedback. For a given trial, the first second corresponded to the ISI, while the target was presented throughout the subsequent five seconds, just as in the training phase. The visual feedback, displayed from 3 s to 6 s, consists of a cursor that starts from the left-middle part of the screen and moves with a fixed velocity to the right part of the screen. The subjects were asked to control the vertical position by modulating their brain activity. During our experiments, the online features (obtained via an autoregressive method) were classified by using the Linear Discriminant Analysis (LDA) method. The present work relies only on the data obtained from the testing phase.

Several neurocognitive questionnaires were proposed to subjects to assess their specific traits such as the self-esteem (3), and the global motivation (4), as well as the ability to perform a motor imagery task (5). Before each session, the subjects' anxiety was also measured (6). To reinforce the learning and to improve the subjects' autonomy, the subjects were asked to train at home and alone with a 10-minute video, which corresponded to 3 training runs without any provided feedback, each day between two sessions. Between two sessions, a new video was sent to the subjects. To obtain accurate EEG head models using surface-based alignment (7), individual T1 sequences (256 sagittal slices, TR = 2.40ms, TE = 2.22ms, 0.80 mm isotropic voxels, $300 \times 320$ matrix; flip angle = 9°) have been obtained by using a 3T Siemens Magnetom PRISMA after the fourth session. The experiments consisted of a 15 minute-resting-state task. Images were preprocessed via the FreeSurfer toolbox (8) and directly imported (15002 vertices) to the Brainstorm toolbox. In this work, we used the Destrieux atlas (9). We digitized the location of the EEG electrodes using the FastTrak 3D digitizer (Polhemus, Inc., VT, USA), the landmarks (nasion, left and right preauricular points), and at the scalp. We aligned these locations with the MRI using the Brainstorm toolbox (10).

**Supplementary Results: Behavioral performance**

Twenty healthy subjects were included in this study. A summary of the demographic information and of the behavioral performances is provided in Table S1. BCI scores correspond to the proportion of hit target. If we observed a significant difference in terms of performances between sessions, it was not the case within a given session. Figure S3 summarizes the results obtained at the run scale.

**Supplementary Results: Spatiotemporal changes**

In this work, we focused our presentation on the $\alpha_2$ and on the $\beta_1$ bands that relied on the use of the IAF. Nevertheless, as a preliminary study, we also performed our analysis within the traditional frequency bands $\theta\,[4-7Hz]$, $\alpha\,[8-12Hz]$, $\beta\,[14-29Hz]$, $\gamma\;[30-40Hz]$. We did not observe a significant difference between the two conditions in the $\theta$ band nor in the $\gamma$ band (Figure S6). Thus, both in terms of diffusion effect and of level of desynchronization, these results did not enable us to observe a clear correlation between the BCI scores and the activations within these bands, contrary to the $\alpha$ and the $\beta$ bands. However, in the latter case, such activations are particularly diffuse (Figure S4 and Figure S5) and more specifically within the $\beta$ band, the statistical results were less robust than those resulting from the use of the IAF (Table S5 and Table S8). We concluded that using restricted frequency bands that rely on the IAF are more selective, and tend to better elicit common trends by taking into account the subjects' specificity.

### Supplementary Figures

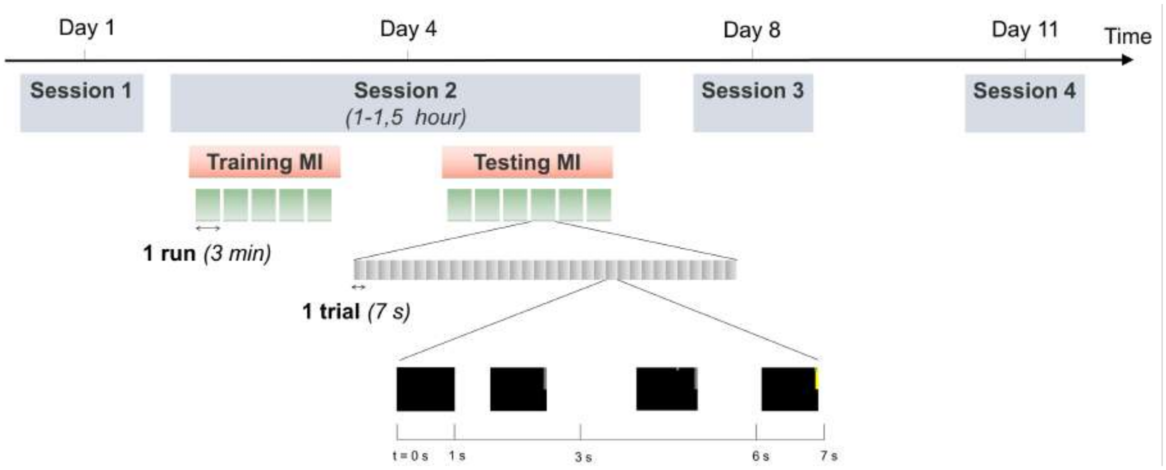


Figure S1: Time course associated with the protocol.

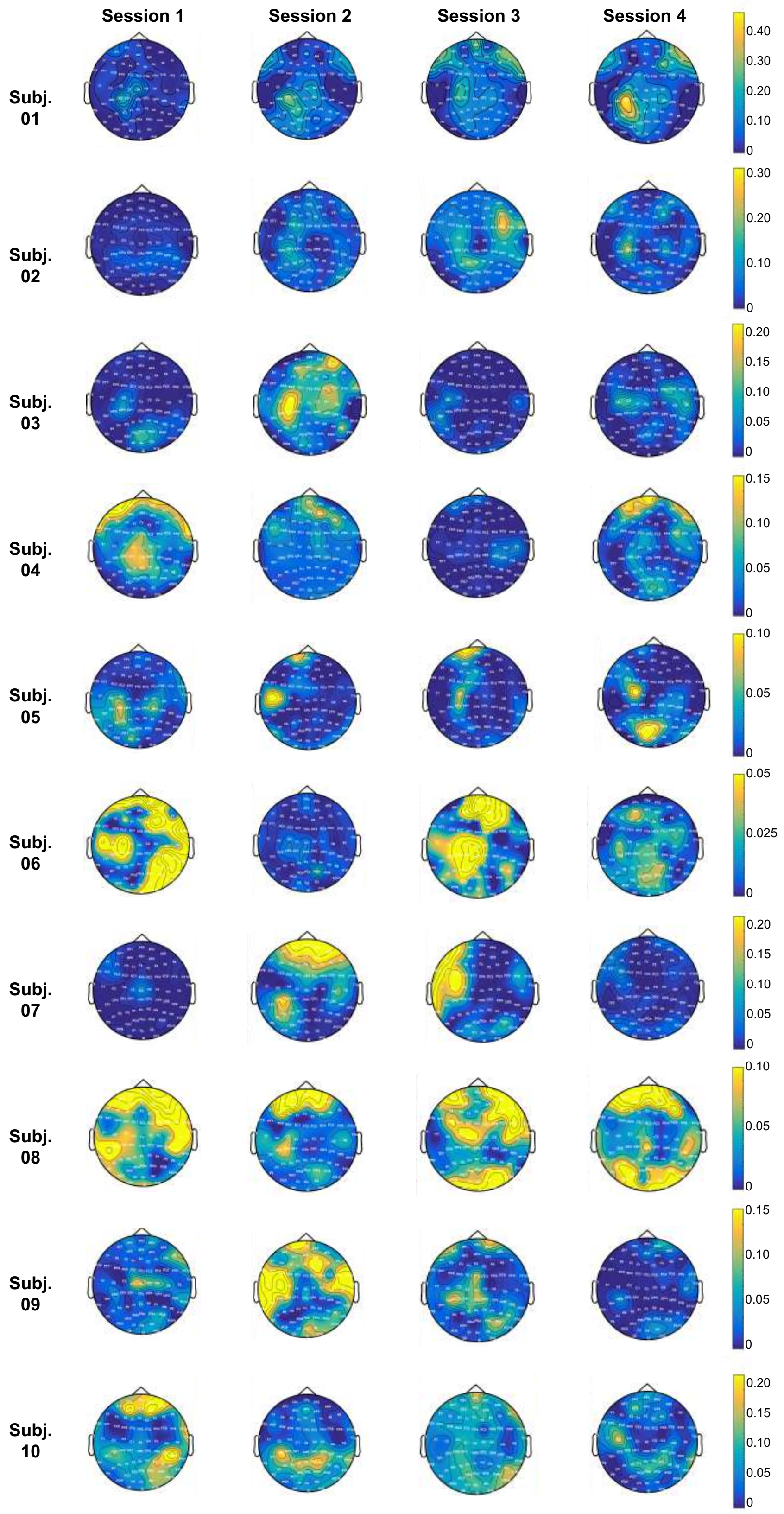


Figure S2: Example of the contrast maps plotted for the first 10 subjects. Contrasts between MI and Rest conditions were computed from data recorded during the training part of the protocol. The colorbars correspond to the R-square value, that gives an estimation of how importantly the means of the two distributions can differ in respect to variance (2).

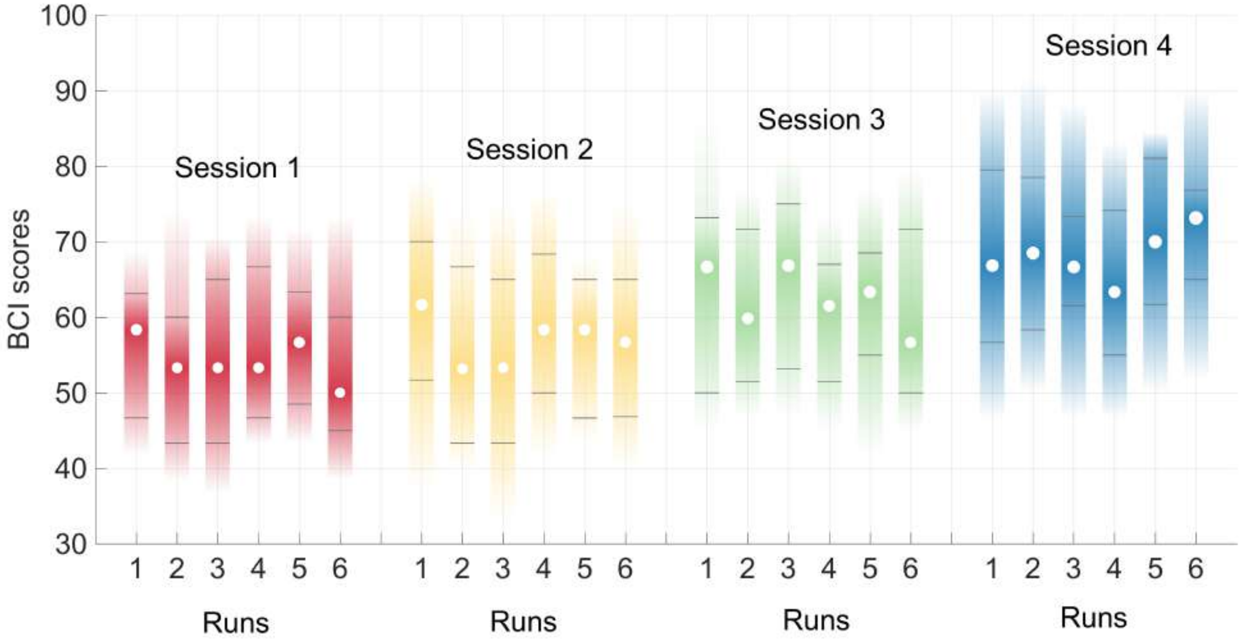


Figure S3: BCI performances obtained across runs. By performing a two-way ANOVA with the session and the runs as within-subject factors, we observed a significant session effect ($F_{3,57} = 13.9$, $p = 6.56.10^{-7}$), and no significant change in BCI performances within a session ($F_{5,95} = 0.26$, $p = 0.935$).

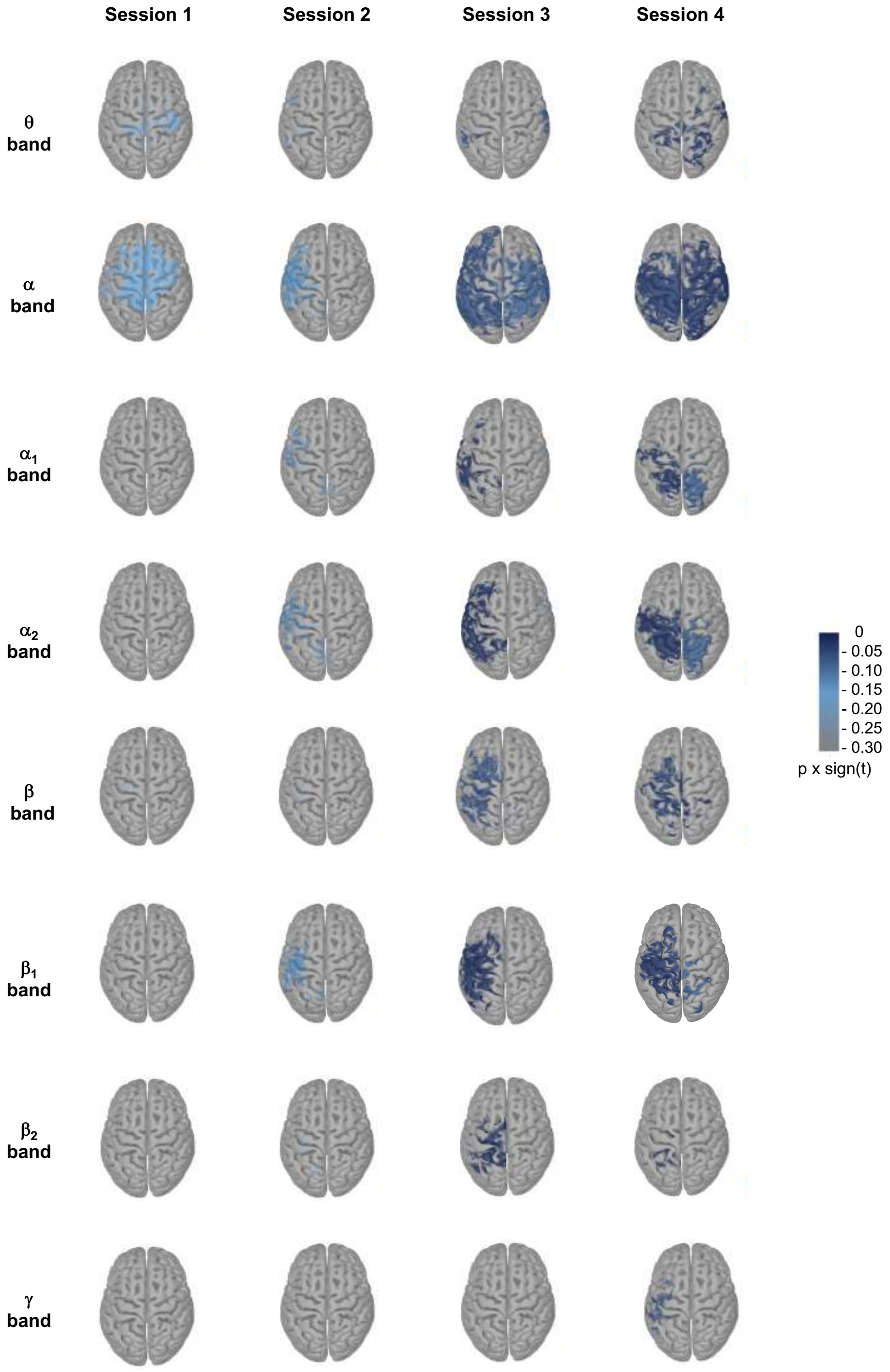


Figure S4: Cluster-based permutation results in all the tested frequency bands computed from the group analysis performed across the 20 subjects within the MNI template (EEG data). Here, we plotted the obtained p-values multiplied by the sign of the t-values resulting from the paired t-test.

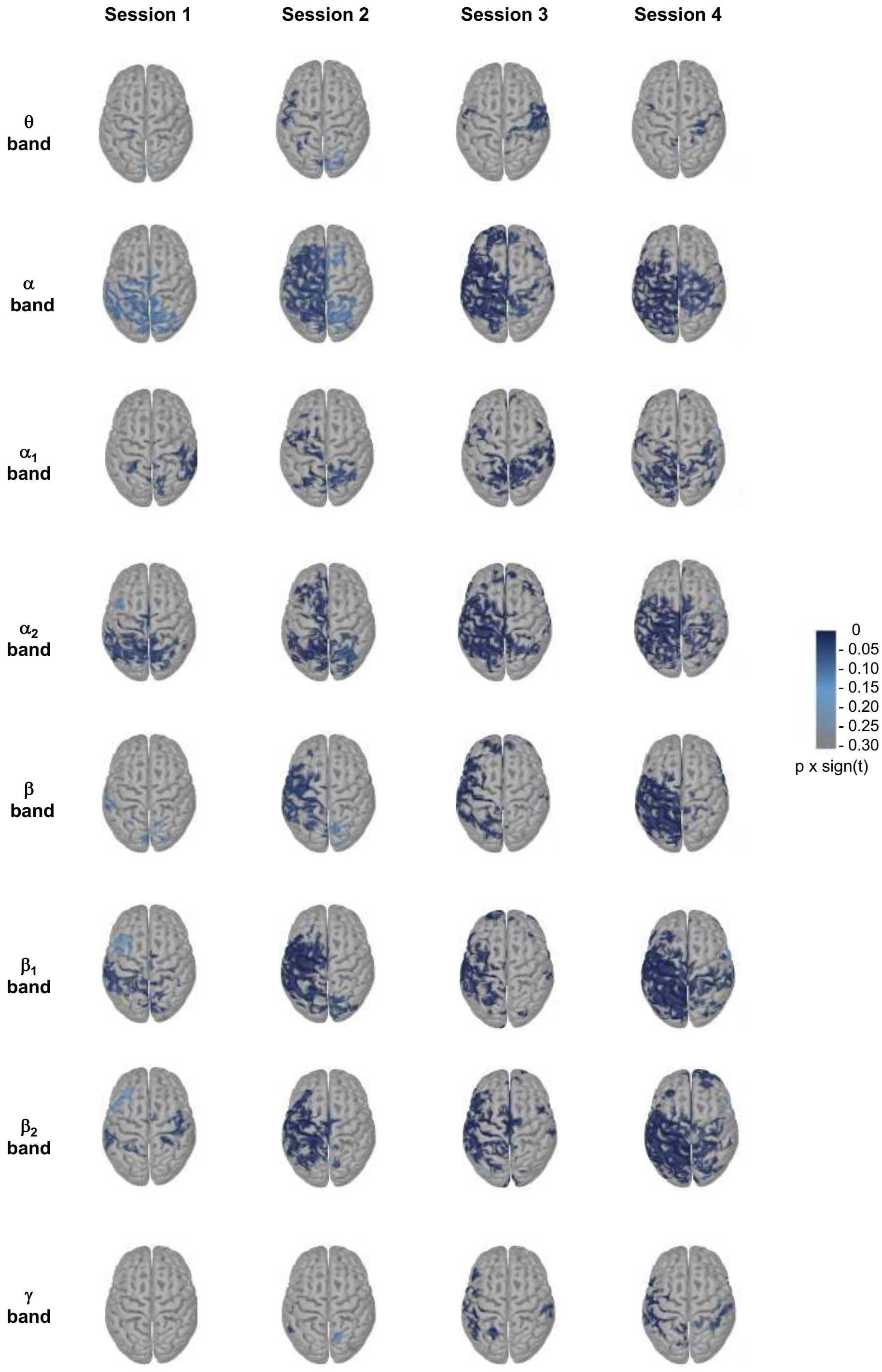


Figure S5: Cluster-based permutation results in all the tested frequency bands computed from the group analysis performed across the 20 subjects within the MNI template (MEG data). Here, we plotted the obtained p-values multiplied by the sign of the t-values resulting from the paired t-test.

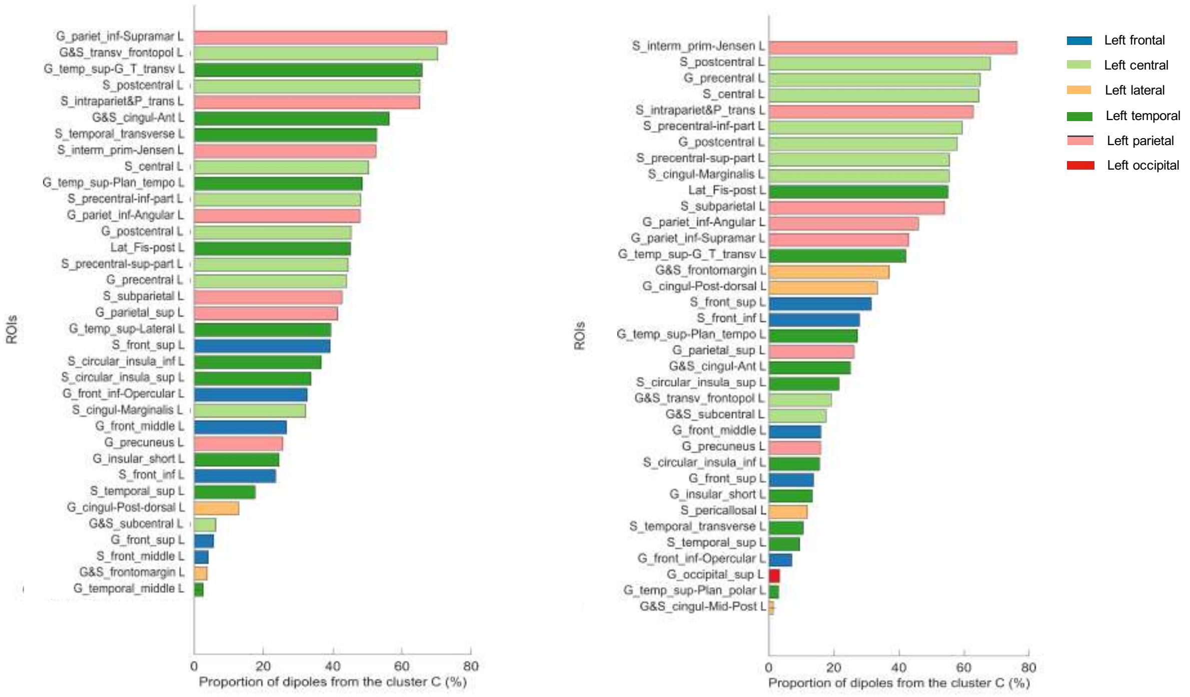


Figure S6: List of the ROIs that appear in the most significant cluster in the $\alpha_2$ (on the left) and the $\beta_1$ band (on the right) during the session 3 (EEG data). Since we expected a desynchronization between the two conditions in terms of activations, we applied a negative one-tailed t-test. Statistics were corrected for multiple comparisons using the cluster approach (11; 10). Cluster-level statistics are obtained by using the sum of the t-values within every cluster. Here, the most significant cluster correspond to the cluster that show the lowest p-value.

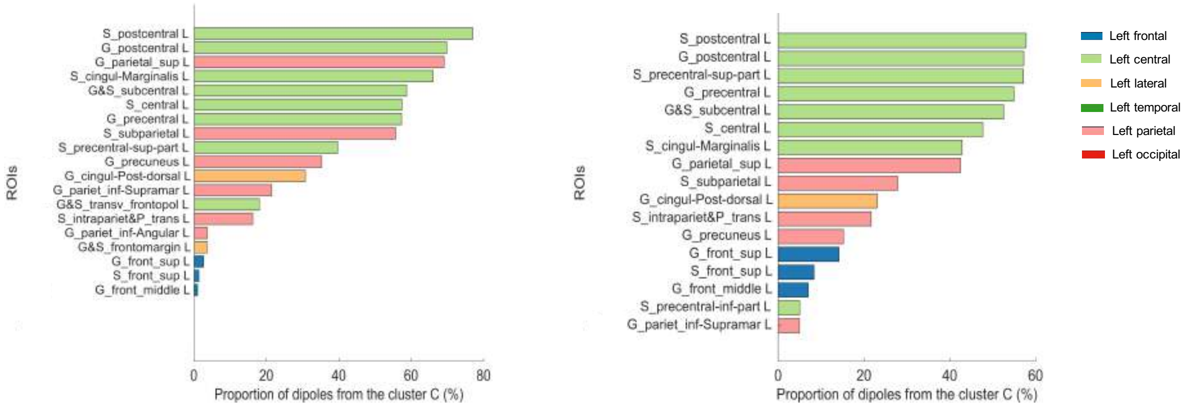


Figure S7: List of the ROIs that appear in the most significant cluster in the $\alpha_2$ (on the left) and the $\beta_1$ band (on the right) during the session 4 (EEG data). Since we expected a desynchronization between the two conditions in terms of activations, we applied a negative one-tailed t-test. Statistics were corrected for multiple comparisons using the cluster approach (11; 10). Cluster-level statistics are obtained by using the sum of the t-values within every cluster. Here, the most significant cluster correspond to the cluster that show the lowest p-value.

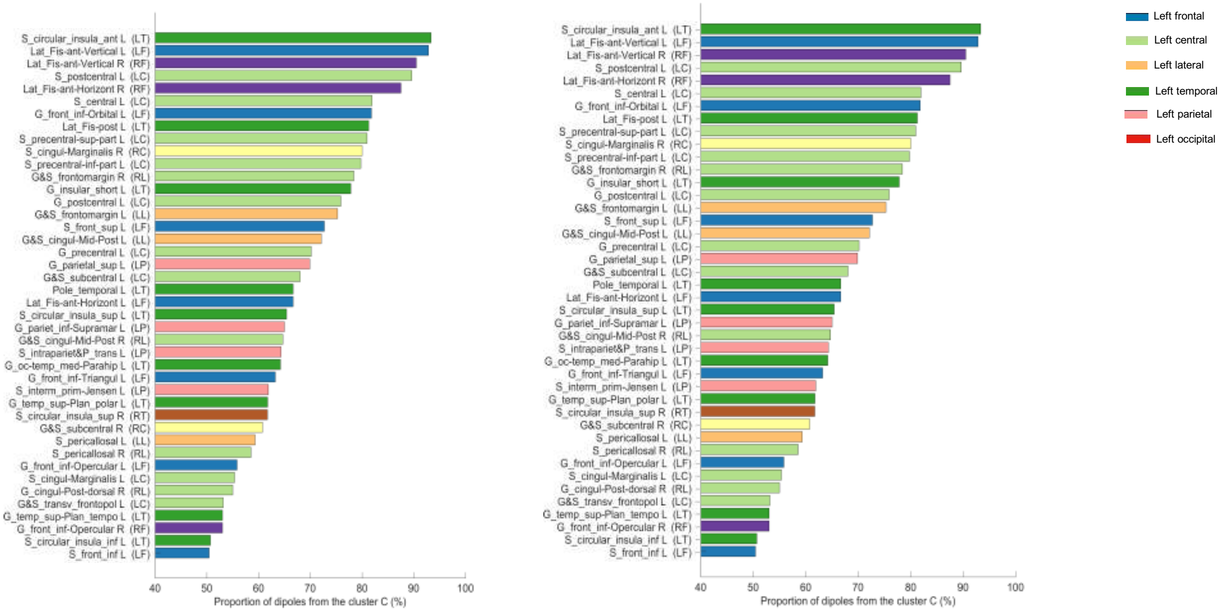


Figure S8: List of the ROIs that appear in the most significant cluster in the $\alpha_2$ (on the left) and the $\beta_1$ band (on the right) during the session 3 (MEG data). Since we expected a desynchronization between the two conditions in terms of activations, we applied a negative one-tailed t-test. Statistics were corrected for multiple comparisons using the cluster approach (11; 10). Cluster-level statistics are obtained by using the sum of the t-values within every cluster. Here, the most significant cluster correspond to the cluster that show the lowest p-value.

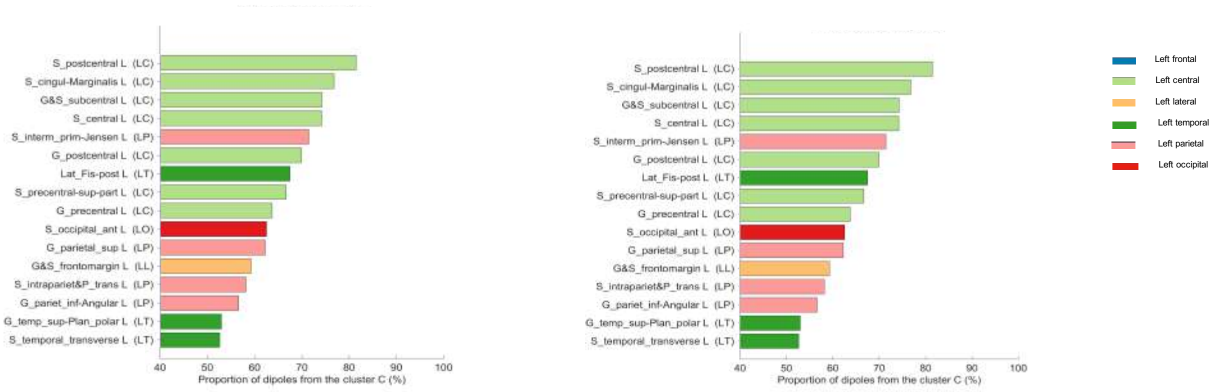


Figure S9: List of the ROIs that appear in the most significant cluster in the $\alpha_2$ (on the left) and the $\beta_1$ band (on the right) during the session 4 (MEG data). Since we expected a desynchronization between the two conditions in terms of activations, we applied a negative one-tailed t-test. Statistics were corrected for multiple comparisons using the cluster approach (11; 10). Cluster-level statistics are obtained by using the sum of the t-values within every cluster. Here, the most significant cluster correspond to the cluster that show the lowest p-value.

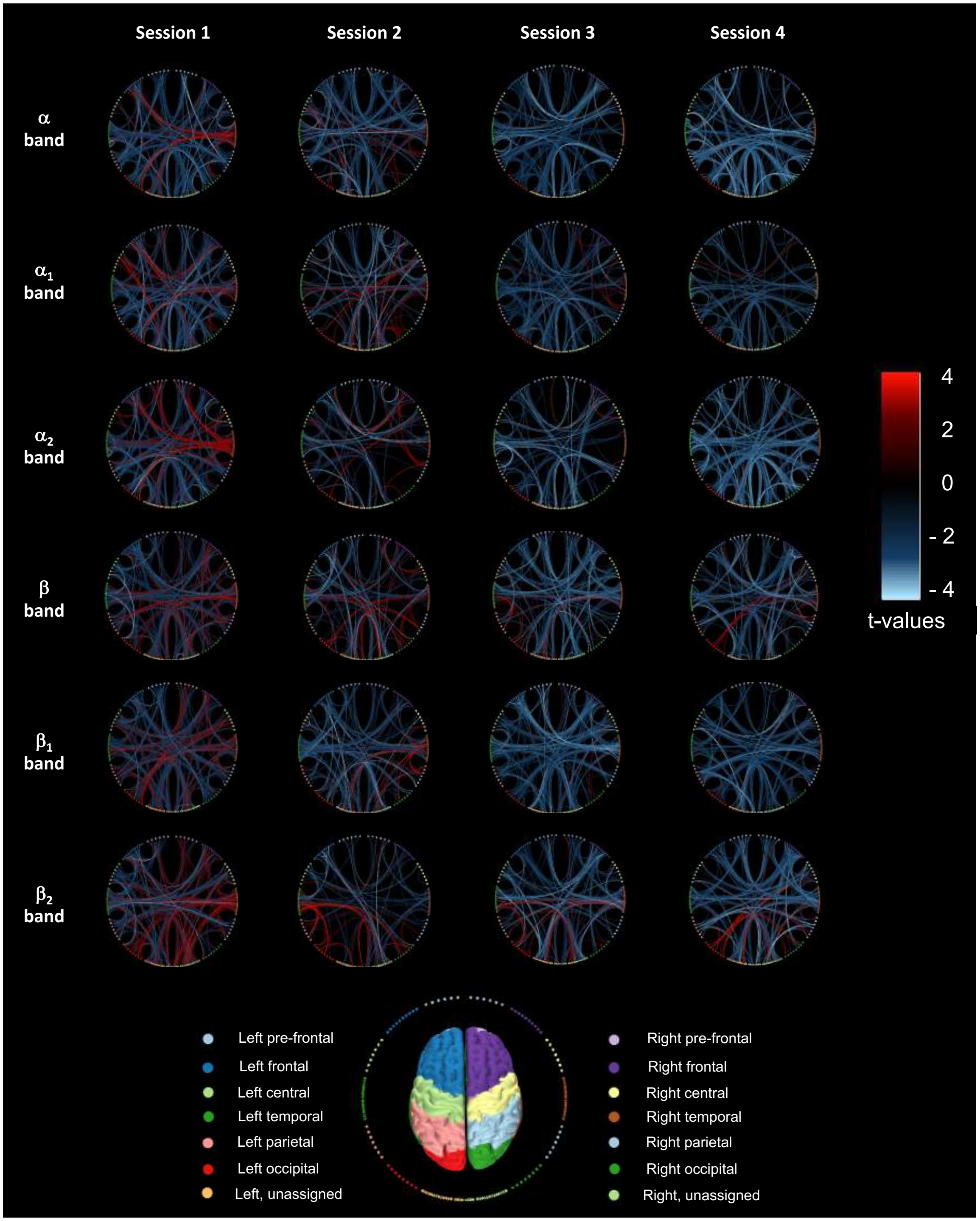


Figure S10: Evolution of the connectomes obtained from two-tailed paired t-tests performed from imaginary coherence values in the $\alpha$ and $\beta$ frequency ranges (EEG data). Here, we showed the contrast between MI and Rest conditions in terms of t-values ($p < 0.005$). Red edges correspond to a higher functional connectivity level in MI condition. Blue edges correspond to a higher functional connectivity level in Rest condition.

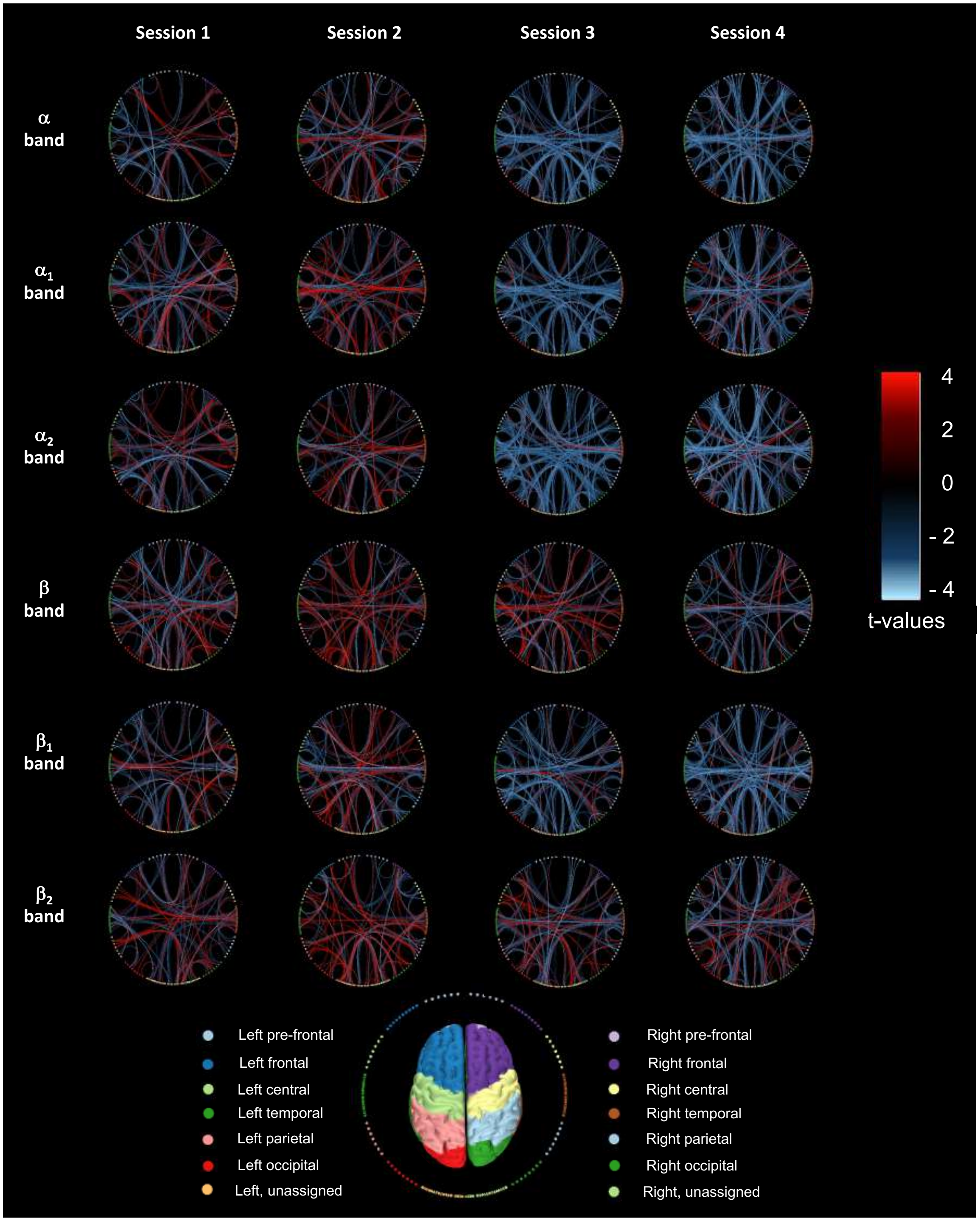


Figure S11: Evolution of the connectomes obtained from two-tailed paired t-tests performed from imaginary coherence values in the $\alpha$ and $\beta$ frequency ranges (MEG data). Here, we showed the contrast between MI and Rest conditions in terms of t-values ($p < 0.005$). Red edges correspond to a higher functional connectivity level in MI condition. Blue edges correspond to a higher functional connectivity level in Rest condition.

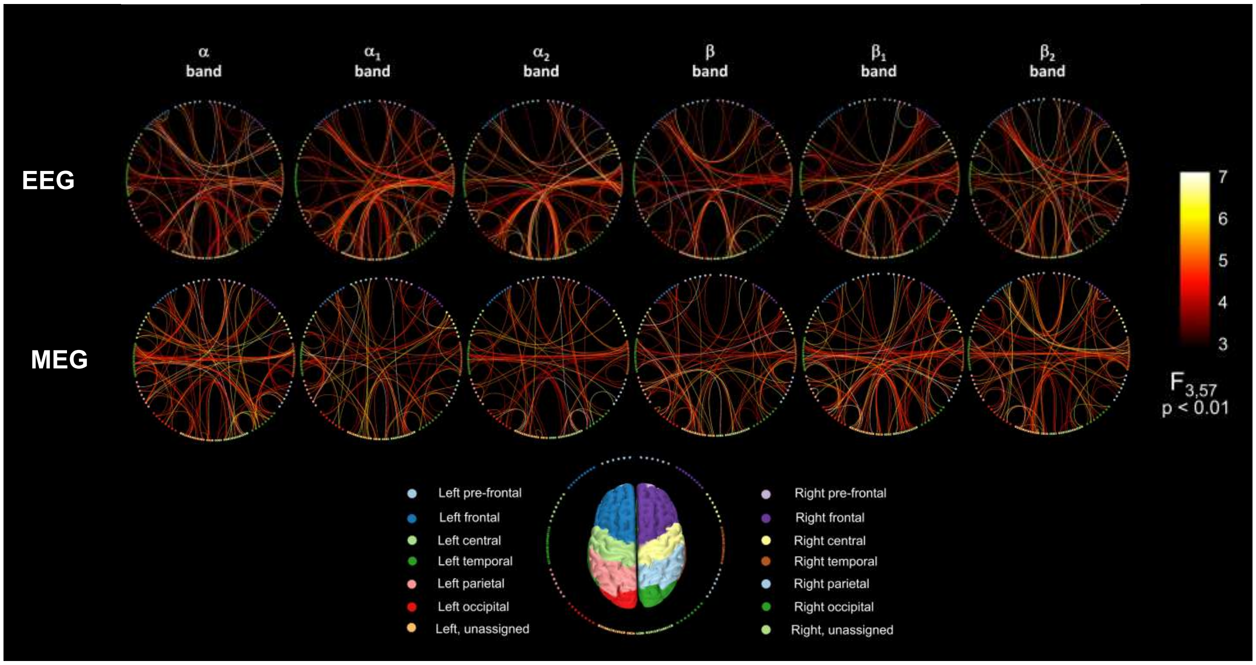


Figure S12: Influence of the training on the functional connectivity. On the top, results from EEG data. On the bottom, results from MEG data. A one-way ANOVA has been computed from the imaginary coherence values, with the session as the within-subject factor ($p < 0.05$). Here, we considered the relative difference between the MI and the rest condition in terms of intensity of connections.

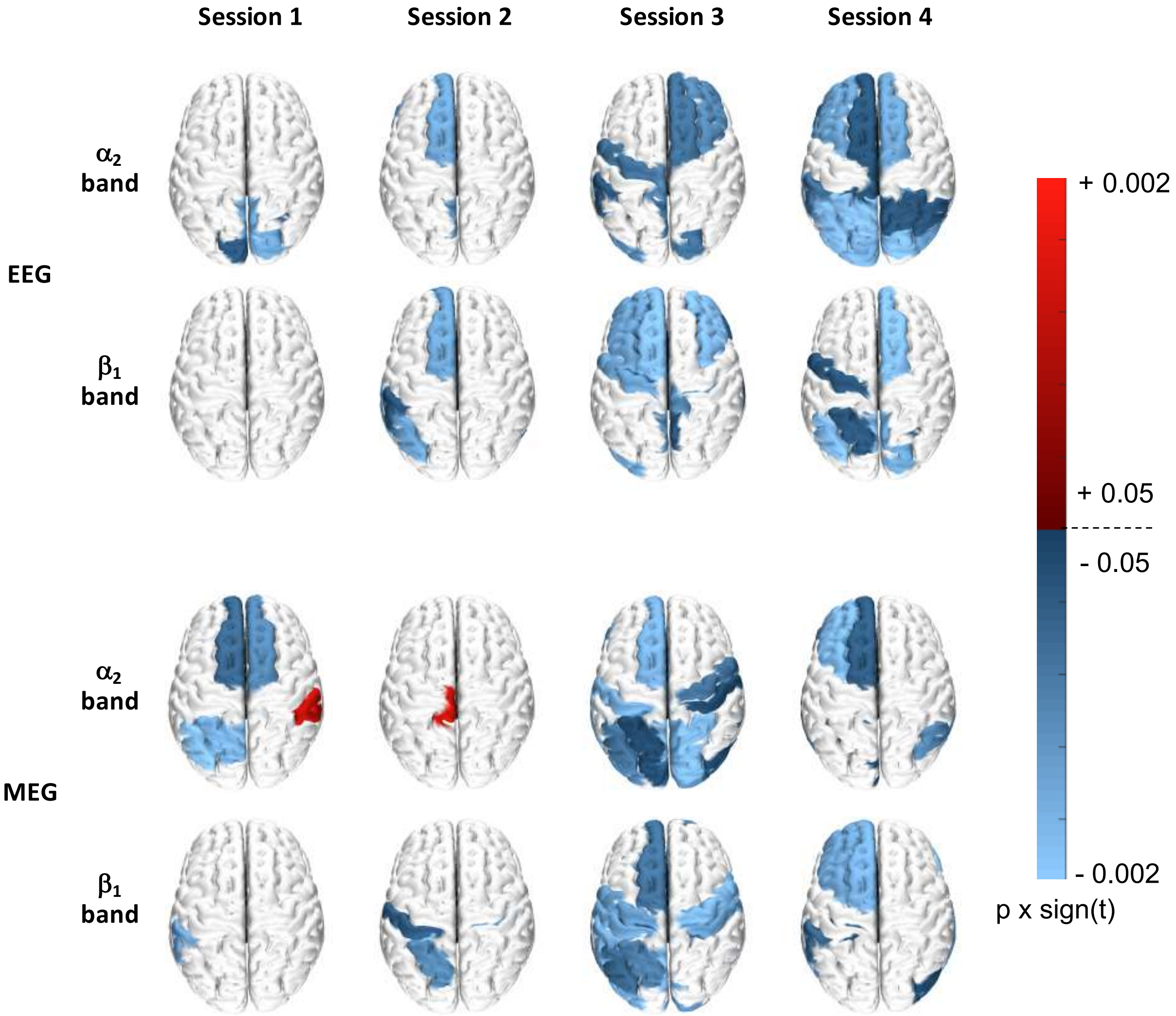


Figure S13: Contrast maps between motor-imagery and rest conditions, in terms of node strength (MEG and EEG data, $p < 0.05$).

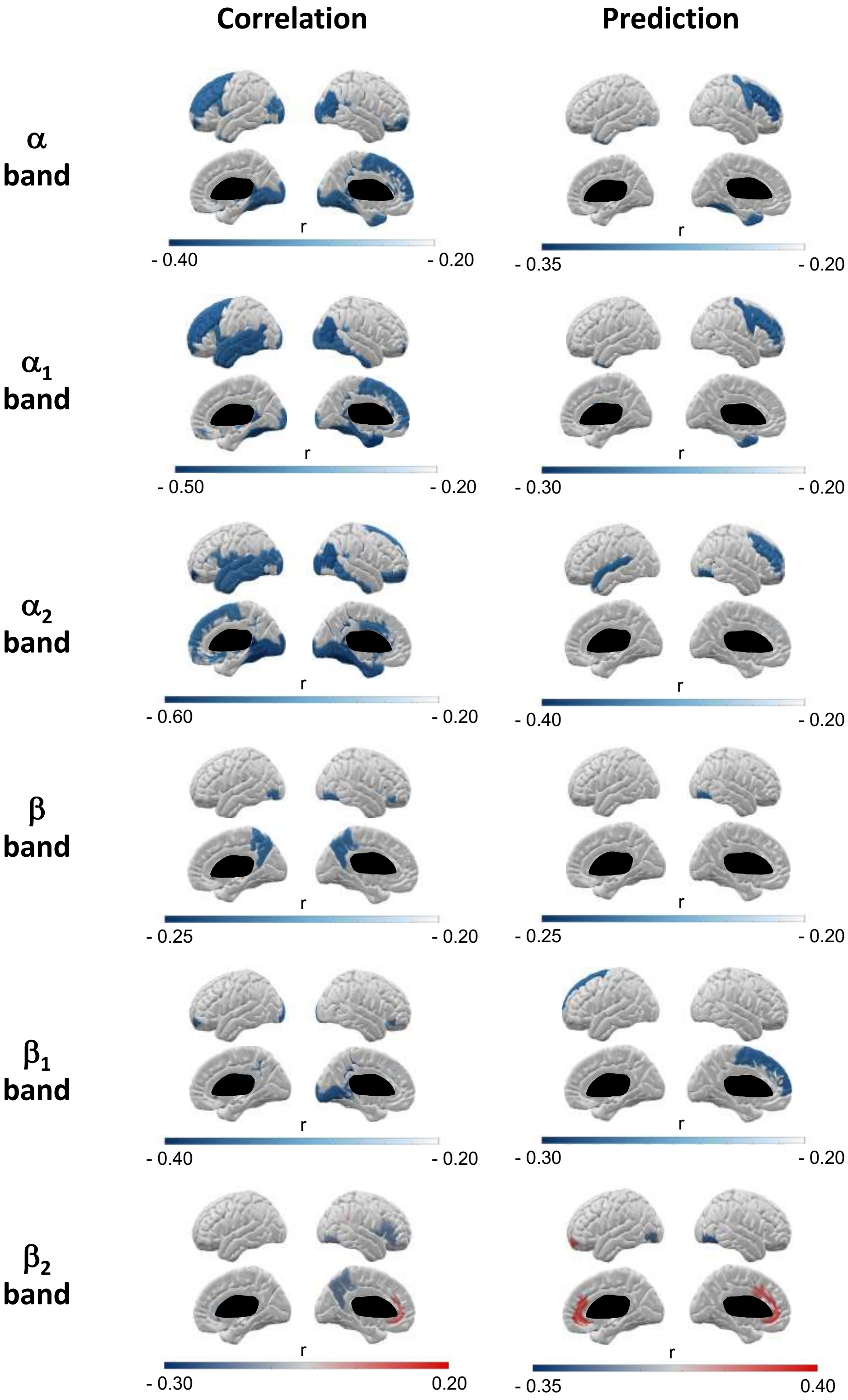


Figure S14: Correlation with BCI performance and prediction of future learning amount with the relative node strength (repeated measures correlation respectively with BCI scores obtained during the same session, and with the learning amount associated with the subsequent session, EEG data, $p < 0.05$).

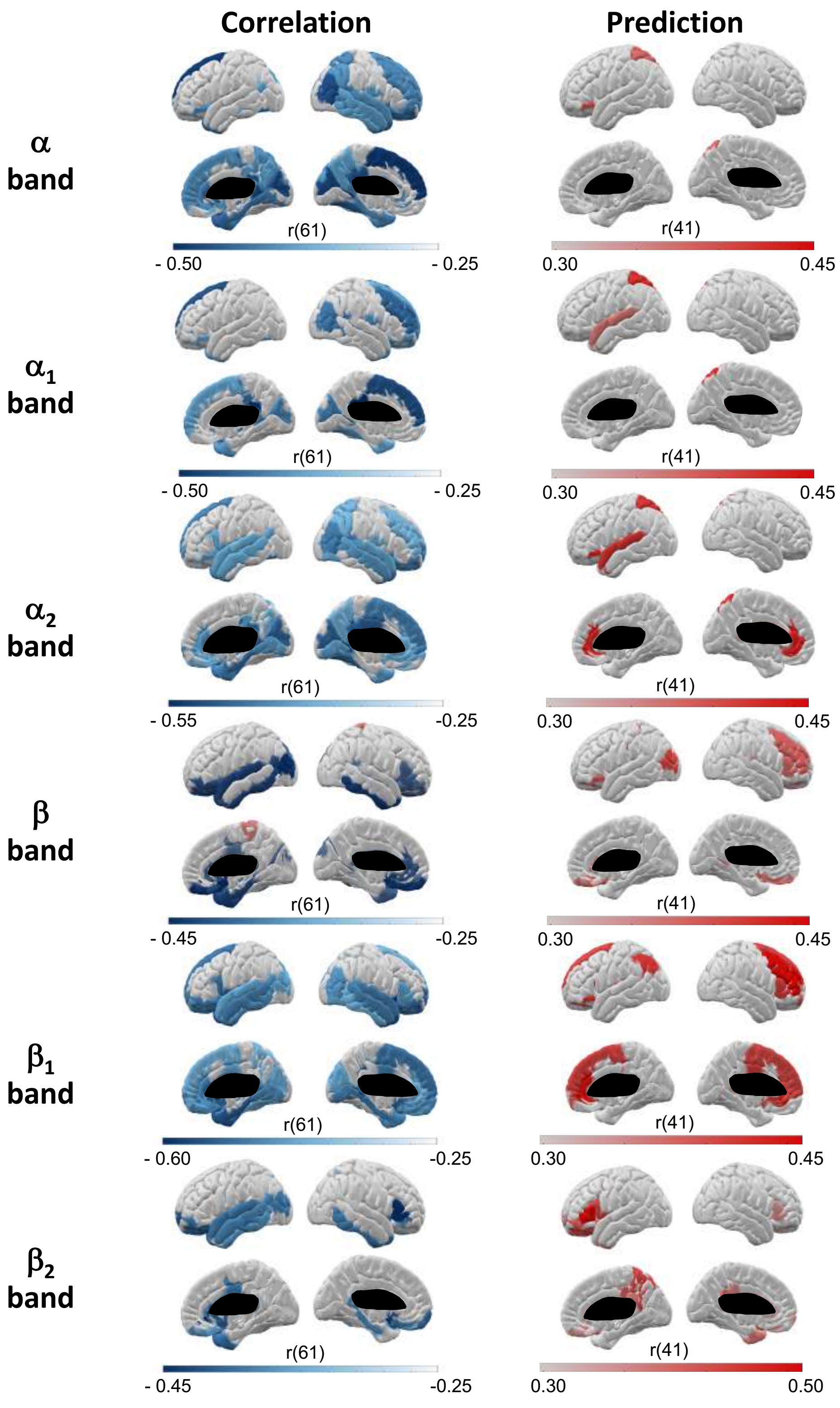


Figure S15: Correlation with BCI performance and prediction of future learning amount with the relative node strength (repeated measures correlation respectively with BCI scores obtained during the same session, and with the learning amount associated with the subsequent session, MEG data, $p < 0.05$).

### Supplementary Tables

Table S1: Summary of participants information. SD stands for standard deviation and Avg stands for average.

| | | | Averaged BCI scores | | | | |
|---|---|---|---|---|---|---|---|
| ID | Age (years) | Gender | Session 1 | Session 2 | Session 3 | Session 4 | **SD** |
| S01 | 19 | Male | 73.8 | 90.2 | 86.6 | 89.3 | 7.61 |
| S02 | 30 | Male | 45.0 | 35.6 | 38.9 | 38.1 | 3.99 |
| S03 | 29 | Male | 66.1 | 61.7 | 72.8 | 86.1 | 10.7 |
| S04 | 31 | Male | 50.0 | 50 | 62.8 | 73.1 | 11.2 |
| S05 | 33 | Male | 51.7 | 51.7 | 61.7 | 63.9 | 6.47 |
| S06 | 26 | Male | 61.2 | 50.6 | 79.3 | 88.9 | 17.3 |
| S07 | 27 | Female | 54.4 | 58.9 | 51.7 | 56.7 | 3.09 |
| S08 | 27 | Female | 59.4 | 66.1 | 56.7 | 70.0 | 6.10 |
| S09 | 22 | Female | 52.8 | 50.0 | 68.9 | 60.0 | 8.44 |
| S10 | 27 | Female | 42.2 | 45.6 | 58.9 | 66.1 | 11.2 |
| S11 | 35 | Female | 53.9 | 45.0 | 55.6 | 63.3 | 7.51 |
| S12 | 29 | Female | 47.2 | 52.2 | 47.8 | 53.9 | 3.29 |
| S13 | 26 | Male | 53.3 | 65.6 | 57.8 | 77.2 | 10.5 |
| S14 | 33 | Female | 54.2 | 50.0 | 60.0 | 69.2 | 8.31 |
| S15 | 22 | Male | 60.0 | 68.9 | 76.1 | 80.0 | 8.80 |
| S16 | 30 | Female | 76.1 | 70.6 | 53.9 | 61.1 | 9.87 |
| S17 | 27 | Male | 48.3 | 61.7 | 66.0 | 71.0 | 9.73 |
| S18 | 28 | Male | 56.0 | 63.9 | 72.0 | 73.0 | 7.94 |
| S19 | 28 | Male | 53.9 | 53.9 | 52.8 | 68.3 | 7.42 |
| S20 | 23 | Male | 45.0 | 57.2 | 58.3 | 56.7 | 6.24 |
| | | **Avg** | 55.2 | 57.5 | 61.9 | 68.3 | 5.76 |
| | | **SD** | 8.95 | 11.8 | 11.4 | 12.6 | |

Table S2: Spearman correlations between demographical or psychological items with, respectively, the BCI performances obtained during the first session and the relative difference between the fourth and the first session in terms of performances. Significant results appear in bold ($p < 0.05$).

| | BCI scores, session 1 | | BCI scores, relative difference | |
|---|---|---|---|---|
| Items | r | p | r | p |
| Age | -0.13 | 0.572 | -0.18 | 0.443 |
| Self-esteem | -0,32 | 0,166 | -0,04 | 0,864 |
| Global motivation | -0.09 | 0.693 | -0.04 | 0.879 |
| Anxiety (Session 1) | -0.15 | 0.526 | 0.04 | 0.862 |
| External visual imagery | 0.26 | 0.271 | -0.04 | 0.864 |
| Internal visual imagery | 0.37 | 0.114 | -0.18 | 0.447 |
| Kinesthetic imagery | **0.45** | **0.045** | -0.20 | 0.41 |

Table S3: Study of the session effect on, respectively, the relative power ($\Delta_P$) and the cluster size ($C_S$). A one-way ANOVA has been computed from the associated values, with the session as the within-subject factor. Significant results appear in bold ($p < 0.03$).

| | EEG | | | | MEG | | | |
|---|---|---|---|---|---|---|---|---|
| | $\Delta_P$ | | CS | | $\Delta_P$ | | CS | |
| Frequency band | $F_{3,57}$ | p | $F_{3,57}$ | p | $F_{3,57}$ | p | $F_{3,57}$ | p |
| $\theta$ | 1.13 | 0.345 | **3.45** | **0.022** | 1.36 | 0.264 | **4.93** | **0.004** |
| $\alpha$ | **5.55** | **0.002** | **6.60** | **6.59.10$^{-4}$** | **4.44** | **0.007** | **4.10** | **0.011** |
| $\alpha_1$ | **5.04** | **0.004** | **5.58** | **0.002** | **3.81** | **0.015** | **3.42** | **0.023** |
| $\alpha_2$ | **6.86** | **5.03.10$^{-4}$** | **5.97** | **0.0013** | **4.18** | **0.010** | **3.37** | **0.025** |
| $\beta$ | 0.42 | 0.735 | **8.81** | **6.84.10$^{-5}$** | **3.88** | **0.014** | **5.82** | **0.002** |
| $\beta_1$ | **6.16** | **0.001** | **6.88** | **4.91.10$^{-4}$** | **4.25** | **0.009** | **3.23** | **0.029** |
| $\beta_2$ | **3.47** | **0.022** | **4.82** | **0.005** | **3.87** | **0.014** | 1.58 | 0.205 |
| $\gamma$ | 0.81 | 0.50 | **4.31** | **0.008** | 0.09 | 0.964 | **4.13** | **0.010** |

Table S4: Study of the session effect on the relative node strength (EEG data). A one-way ANOVA has been computed from the $\Delta_N$ values, with the session as the within-subject factor. Here, we only show the significant results ($p < 0.025$).

| Frequency bands | ROI | Associated region | $F_{3,57}$ | p |
|---|---|---|---|---|
| $\alpha$ band | Cuneus R | Occipital R | 3.70 | 0.017 |
| | Lingual gyrus R | Occipital L | 2.78 | 0.049 |
| | Superior occipital gyrus R | Occipital R | 3.98 | 0.012 |
| | Middle occipital sulcus and lunatus sulcus L | | 4.30 | 0.008 |
| | Superior occipital sulcus and transverse occipital sulcus L | | 4.00 | 0.012 |
| | Parieto-occipital sulcus R | | 3.38 | 0.024 |
| $\alpha_1$ band | Superior occipital gyrus R | Occipital R | 4.06 | 0.011 |
| | Posterior transverse collateral sulcus R | Occipital R | 3.98 | 0.012 |
| | Inferior temporal gyrus R | Temporal R | 3.85 | 0.014 |
| | Superior temporal sulcus R | Temporal R | 3.83 | 0.014 |
| | Cuneus R | Occipital R | 3.80 | 0.015 |
| | Middle occipital gyrus R | Occipital R | 3.71 | 0.016 |
| | Vertical ramus of the anterior segment of the lateral sulcus L | Frontal L | 3.61 | 0.019 |
| | Posterior transverse collateral sulcus L | Occipital L | 3.57 | 0.020 |
| | Calcarine sulcus L | Occipital L | 3.56 | 0.020 |
| | Pole occipital L | Occipital L | 3.42 | 0.023 |
| | Superior occipital sulcus and transverse occipital sulcus R | | 3.39 | 0.024 |
| $\alpha_2$ band | Middle occipital sulcus and lunatus sulcus L | | 5.98 | 0.0013 |
| | Cuneus R | Occipital R | 5.71 | 0.0017 |
| | Subparietal sulcus R | Parietal R | 4.72 | 0.005 |
| | Subparietal sulcus L | Parietal L | 4.34 | 0.008 |
| | Middle occipital gyrus R | Occipital R | 3.99 | 0.012 |
| | Superior occipital sulcus and transverse occipital sulcus L | | 3.94 | 0.013 |
| | Parieto-occipital sulcus R | | 3.94 | 0.013 |
| | Horizontal ramus of the anterior segment of the lateral sulcus L | Frontal L | 3.81 | 0.015 |
| | Superior occipital gyrus R | Occipital R | 3.80 | 0.015 |
| | Posterior ramus of the lateral sulcus R | Temporal R | 3.72 | 0.016 |
| | Cuneus L | Occipital L | 3.71 | 0.017 |
| | Middle occipital gyrus L | Occipital L | 3.60 | 0.019 |
| | Posterior-ventral part of the cingulate gyrus R | Lateral R | 3.56 | 0.020 |
| | Superior temporal sulcus R | Temporal R | 3.42 | 0.023 |
| | Posterior transverse collateral sulcus L | Occipital L | 3.41 | 0.024 |
| | Middle occipital sulcus and lunatus sulcus R | | 3.38 | 0.024 |
| $\beta$ band | Paracentral lobule and sulcus L | | 4.56 | 0.006 |
| | Transverse frontopolar gyri and sulci L | | 3.60 | 0.019 |
| | Subparietal sulcus L | Parietal L | 4.65 | 0.006 |
| $\beta_1$ band | Subparietal sulcus L | Parietal L | 6.88 | $4.92.10^{-4}$ |
| | Subparietal sulcus R | Parietal R | 4.15 | 0.01 |
| | Marginal branch (or part) of the cingulate sulcus L | Central L | 3.74 | 0.016 |
| | Paracentral lobule and sulcus L | | 3.53 | 0.020 |
| $\beta_2$ band | Transverse frontopolar gyri and sulci L | | 4.20 | 0.01 |
| | Triangular part of the inferior frontal gyrus R | Frontal R | 4.20 | 0.01 |
| | Fronto-marginal gyrus R | | 4.01 | 0.012 |
| | Paracentral lobule and sulcus L | | 3.73 | 0.016 |
| | Inferior frontal sulcus R | Frontal R | 3.65 | 0.018 |
| | Anterior segment of the circular sulcus of the insula R | Temporal R | 3.14 | 0.030 |

Table S5: Study of the session effect on the relative node strength (MEG data). A one-way ANOVA has been computed from the $\Delta_N$ values, with the session as the within-subject factor. Here, we only show the significant results ($p < 0.025$).

| Frequency bands | ROI | Associated region | $F_{3,57}$ | p |
|---|---|---|---|---|
| $\alpha$ band | Middle occipital gyrus R | Occipital R | 5.16 | 0.003 |
| | Cuneus R | Occipital R | 4.91 | 0.004 |
| | Orbital part of the inferior frontal gyrus R | Frontal R | 4.76 | 0.005 |
| | Superior occipital gyrus R | Occipital R | 4.39 | 0.008 |
| | Anterior part of the cingulate gyrus and sulcus (ACC) R | | 4.20 | 0.009 |
| | Posterior-ventral part of the cingulate gyrus (vPCC) R | Lateral R | 3.94 | 0.013 |
| | Occipital pole R | Occipital R | 3.81 | 0.015 |
| | Anterior part of the cingulate gyrus and sulcus (ACC) L | | 3.63 | 0.018 |
| | Inferior temporal gyrus L | Temporal L | 3.50 | 0.021 |
| | Temporal pole R | Temporal R | 3.42 | 0.023 |
| | Cuneus L | Occipital L | 3.37 | 0.024 |
| $\alpha_1$ band | Short insular gyri L | Temporal L | 5.26 | 0.003 |
| | Superior occipital gyrus R | Occipital R | 4.44 | 0.007 |
| | Cuneus R | Occipital R | 4.19 | 0.009 |
| | Lateral aspect of the superior temporal gyrus L | Temporal L | 3.60 | 0.019 |
| | Superior occipital gyrus L | Occipital L | 3.36 | 0.024 |
| $\alpha_2$ band | Orbital gyri R | Prefrontal R | 4.35 | 0.008 |
| | Occipital pole R | Occipital R | 4.32 | 0.008 |
| | Cuneus R | Occipital R | 3.81 | 0.015 |
| | Cuneus L | Occipital L | 3.80 | 0.015 |
| | Anterior part of the cingulate gyrus and sulcus (ACC) R | | 3.60 | 0.019 |
| | Middle occipital gyrus R | Occipital R | 3.54 | 0.020 |
| | Anterior part of the cingulate gyrus and sulcus (ACC) L | | 3.54 | 0.020 |
| | Anterior transverse temporal gyrus L | Temporal L | 3.46 | 0.022 |
| | Lingual gyrus L | Occipital L | 3.46 | 0.022 |
| | Posterior-ventral part of the cingulate gyrus (vPCC) R | Lateral R | 3.41 | 0.023 |
| $\beta$ band | N.A. | N.A. | N.A. | N.A. |
| $\beta_1$ band | Posterior-ventral part of the cingulate gyrus (vPCC) R | Lateral R | 4.09 | 0.011 |
| | Short insular gyri R | Temporal R | 4.02 | 0.011 |
| | Parahippocampal gyrus R | Temporal R | 3.96 | 0.012 |
| | Orbital part of the inferior frontal gyrus R | Frontal R | 3.75 | 0.016 |
| | Anterior part of the cingulate gyrus and sulcus (ACC) L | | 3.71 | 0.017 |
| | Orbital gyri R | Prefrontal R | 3.59 | 0.020 |
| | Parahippocampal gyrus L | Temporal L | 3.49 | 0.021 |
| | Subcallosal gyrus L | Lateral L | 3.42 | 0.023 |
| $\beta_2$ band | Temporal pole R | Temporal R | 4.94 | 0.004 |
| | Anterior transverse temporal gyrus R | Temporal R | 4.45 | 0.007 |

Table S6: Correlation with BCI performance and prediction of futures scores with, respectively, the relative power ($\Delta_P$) between the two conditions, and the cluster size (CS). The most significant results (from repeated measures correlations (12)) appear in bold ($p < 0.006$).

| | EEG | | | | | | | | MEG | | | | | | | |
|---|---|---|---|---|---|---|---|---|---|---|---|---|---|---|---|---|
| | Correlation | | | | Prediction | | | | Correlation | | | | Prediction | | | |
| | $\Delta_P$ | | CS | | $\Delta_P$ | | CS | | $\Delta_P$ | | CS | | $\Delta_P$ | | CS | |
| Frequency band | r | p | r | p | r | p | r | p | r | p | r | p | r | p | r | p |
| $\theta$ | -0.13 | 0.33 | **0.45** | **3.0.10$^{-4}$** | 0.04 | 0.79 | 0.005 | 0.98 | -0.12 | 0.365 | **0.44** | **3.0.10$^{-4}$** | 0.03 | 0.87 | -0.06 | 0.69 |
| $\alpha$ | **-0.37** | **0.003** | **0.56** | **3.10.10$^{-6}$** | 0.04 | 0.79 | -0.15 | 0.34 | -0.27 | 0.035 | **0.39** | **0.002** | 0.12 | 0.45 | -0.14 | 0.37 |
| $\alpha_1$ | **-0.48** | **1.0.10$^{-4}$** | **0.42** | **7.0.10$^{-4}$** | 0.15 | 0.36 | -0.05 | 0.77 | **-0.38** | **0.003** | **0.36** | **0.005** | 0.20 | 0.21 | -0.13 | 0.40 |
| $\alpha_2$ | **-0.63** | **6.30.10$^{-8}$** | **0.57** | **1.50.10$^{-6}$** | 0.15 | 0.34 | -0.18 | 0.26 | **-0.54** | **7.61.10$^{-6}$** | **0.53** | **1.04.10$^{-5}$** | 0.23 | 0.16 | -0.21 | 0.18 |
| $\beta$ | 0.03 | 0.81 | **0.48** | **1.0.10$^{-4}$** | -0.10 | 0.53 | -0.01 | 0.95 | -0.29 | 0.023 | **0.36** | **0.004** | -0.01 | 0.97 | 0.02 | 0.90 |
| $\beta_1$ | **-0.57** | **1.5.10$^{-6}$** | **0.60** | **2.7.10$^{-7}$** | 0.11 | 0.50 | -0.18 | 0.29 | **-0.44** | **3.0.10$^{-4}$** | **0.46** | **1.0.10$^{-4}$** | 0.14 | 0.38 | -0.09 | 0.58 |
| $\beta_2$ | -0.33 | 0.01 | **0.42** | **7.0.10$^{-4}$** | -0.01 | 0.95 | -0.06 | 0.73 | -0.28 | 0.027 | **0.36** | **0.004** | -0.13 | 0.43 | 0.04 | 0.8 |
| $\gamma$ | -0.17 | 0.20 | **0.47** | **1.0.10$^{-4}$** | -0.04 | 0.78 | -0.08 | 0.63 | 0.02 | 0.877 | 0.32 | 0.012 | -0.02 | 0.86 | 0.15 | 0.33 |
